# A Continuously Oxygenated Macroencapsulation System Enables High-Density Packing and Delivery of Insulin-Secreting Cells

**DOI:** 10.1101/2025.04.21.649806

**Authors:** Tung T. Pham, Phuong L. Tran, Linda A. Tempelman, Simon G. Stone, Christopher Piccirillo, Alan Li, James A. Flanders, Minglin Ma

## Abstract

The encapsulation of insulin-secreting cells within immuno-protective systems holds significant promise for curative treatment of type 1 diabetes without immunosuppression. A major challenge, however, remains the inadequate oxygen tension within the encapsulation systems, which compromises the survival and function of encapsulated cells and necessitates low packing density and impractically large systems to deliver a curative cell mass. In this study, we present a novel cell encapsulation system capable of generating oxygen via the electrolysis of tissue moisture to provide a continuous oxygen supply to densely packed insulin-secreting cells. Our system comprises a miniaturized implantable electrochemical oxygen generator (iEOG) and a scalable cylindrical cell encapsulation pouch, designed in a linear configuration to facilitate minimally invasive implantation and retrieval. The oxygen generation from the system was shown to be precisely controlled, stable, and capable of supporting clinically relevant doses of pancreatic islets. *In vitro* studies demonstrated that the oxygenated system effectively maintained the viability and function of insulinoma cell aggregates and human pancreatic islets at densities of 60,000 islet equivalents per mL (IEQ/mL) or 4,200 IEQ/cm² under a hypoxic cell culture condition (1% O ). In an allogeneic rat model, the oxygenated systems containing pancreatic islets implanted into the poorly vascularized but clinically attractive subcutaneous space at a density of 60,000 IEQ/mL successfully reversed diabetes for up to about 3 months without the need for immunosuppression, while animals implanted with non-oxygenated systems remained diabetic. Most (∼ 90%) of the pancreatic islets encapsulated in the continuously oxygenated systems were found viable and functional upon retrieval. These findings suggest the feasibility of using continuous oxygenation to support insulin-secreting cells at high loading densities in subcutaneous space, enabling the development of an encapsulation system with clinically practical dimensions.

## Introduction

Type 1 diabetes (T1D) is a chronic disorder caused by the autoimmune destruction of pancreatic β-cells, which eventually leads to insulin deficiency. Exogenous insulin therapy, the current mainstay to manage this disease, necessitates a stringent and constant effort from patients to carefully manage insulin intake balanced with diet, exercise, sleep, and stress. Despite the significant advances in insulin therapy, it remains non-curative and incapable of maintaining glucose homeostasis, predisposing patients to life-threatening complications including severe hypoglycemia events (SHEs) [1–3] and irreversible organ damage caused by chronic hyperglycemia [4]. Cellular therapy has emerged as a potential cure to overcome this challenge given the ability of primary and stem cell-derived insulin-producing cells to inherently respond to blood glucose changes. Multicenter clinical trials of intraportal transplantation of allogenic cadaveric islets in over 1200 patients have consistently demonstrated the superior efficacy and safety of cell therapy, evidenced by physiological glycemic control without exogenous insulin and nearly complete eradication of SHEs [5, 6]. Favorable outcomes of the recent Phase 1/2 clinical trial by Vertex Pharmaceuticals (NCT04786262) further confirmed the feasibility of cell replacement therapy for T1D with the limitation of donor availability being addressed by the sustainable supply of stem cell-derived β cells (SC-β cells). However, the requisite lifelong immunosuppression with deleterious side effects, such as nephrotoxicity, opportunistic infections, and malignancies, remains a formidable obstacle to the widespread clinical application of current cellular therapies [7]. Thus, developing an immunosuppression-free strategy to protect the transplanted cells is imperative to fully harness the potential of cell therapy for curing T1D.

Over the past few decades, the concept of cell encapsulation has been proposed to circumvent immunosuppression. In principle, enveloping insulin-secreting cells within a semipermeable system could prevent direct contact with the detrimental elements of the immune system while allowing for efficient diffusion of glucose, insulin, nutrients, and metabolic wastes. Among encapsulation strategies, macroencapsulation recently garnered more attraction in the field, especially after SC-β cells became clinically available. As opposed to microencapsulation, cell confinement and the ability to retrieve the whole system of the macroencapsulation devices can address safety and regulatory concerns. Furthermore, subcutaneous space stands out as an ideal transplantation site given the large capacity, minimally invasive surgical procedure, amenability to frequent monitoring, and safe retrievability. However, despite notable insights from research endeavors, translating this concept into clinical applications has been impeded by persistent and intertwined challenges including insufficient supply of oxygen, foreign body responses (FBR), and inability of maintaining a curative dose of cells functional in a reasonably sized system. Particularly, β-cells have an intrinsically high oxygen consumption rate due to the high metabolic demand for insulin synthesis and secretion. Native pancreatic islets require a disproportionately high proportion of pancreatic blood flow relative to their mass [8, 9]. However, the oxygen supply in encapsulation systems is inadequate due to the lack of direct vascularization of encapsulated cells and relatively low oxygen tension in extrahepatic sites (such as the desirable subcutaneous milieu), leading to hypoxia-induced cell death [10–12] and irreversibly impaired β-cell function [13, 14]. The oxygen deficit is further exacerbated by the low oxygen permeability of fibrotic tissues surrounding the implants caused by foreign body response against the encapsulating materials [15]. Moreover, insufficient oxygen supply also limits the packing density of β-cells within the encapsulation system as escalating cell density amplifies the requirement for oxygen owing to the steeper oxygen gradient among cell layers [16–20]. As a result, low cell densities are typically used to reduce hypoxia-induced cell loss post-implantation, rendering the requisite implant dimensions impractical for delivering a curative dose of β-cells [8, 21]. This limitation in packing density has not always been noted in small animal studies but poses a significant barrier to scaling up for clinical translation.

Enhanced oxygen supply to cell encapsulation systems could potentially address the aforementioned challenges. Integration of oxygen-generating biomaterials (e.g. calcium peroxide [16, 18, 22], sodium percarbonate [23, 24], and lithium peroxide [25]) into the encapsulation systems has shown encouraging outcomes in preclinical models. The common downsides of these chemical-based approaches are the short duration of oxygen supply, uncontrolled oxygen generation rate, and undesirable end products [16, 19]. Another approach, by BetaO2 Technologies, tackled these challenges by integrating a refillable oxygen reservoir into a macroencapsulation system, allowing for oxygen replenishment via percutaneous injection ports. Devices supplied daily with supraphysiological levels of oxygen demonstrated long-term diabetes reversal in both an allogeneic rat-to-rat and a xenogeneic rat-to-pig model [26–28]. Nonetheless, the requirement for oxygen replenishing via frequent injections and device bulkiness pose ongoing challenges.

To address this collection of issues, we report here a BioElectronics-Assisted Macroencapsulation (BEAM) system capable of providing a continuous oxygen supply to encapsulated cells via electrolysis of tissue moisture. The components of the implantable electrochemical oxygen generator (iEOG) are assembled in a compact, enclosed design (13 mm diameter x 3.1 mm thick), which is then connected to a linear core-shell cell pouch. Islets are housed within the annular space between an inner gas-permeable silicon tubing and an outer immunoprotective tubing made of a hydrogel-impregnated electrospun membrane. This unique design allows for optimal mass transportation and proximity between cells and the oxygen source. The concentric configuration augments the surface-to-volume ratio and facilitates scalability both longitudinally and radially. The edge-free, round-end cylindrical design of the cell encapsulation pouch eliminates the need for intricate sealing that could lead to sharp edges and rigid connections, triggering tissue irritation and fibrosis. The final linear layout of the system simplifies implantation and retrieval through minor incisions using minimally invasive techniques.

We demonstrated consistent and programmable oxygen generation from the system in both benchtop and *in vivo* contexts, enabling adaptability to various cell types and cell dosages. The beneficial impacts of oxygenation on the viability and secretion of insulin were demonstrated *in vitro* on insulinoma-1 cell (INS-1) aggregates and human pancreatic islets encapsulated at a high density (60,000 IEQ/mL). Additionally, we evaluated the therapeutic potential of the BEAM system after subcutaneous implantation in an allogeneic diabetic rat model and achieved a ∼3-month cure and up to 90% cell viability in retrieved devices. Overall, the BEAM system provides a feasible approach to preserving the viability and function of densely packed insulin-secreting cells in the poorly vascularized but clinically attractive subcutaneous site, thus paving the way for development of a practical cell encapsulation system with realistic dimensions and minimally invasive procedures for immunosuppression-free T1D cell replacement therapy.

### Design and Fabrication of the BEAM system

The BEAM system comprises two integral components: an implantable electrochemical oxygen generator engineered to produce a continuous and controllable supply of oxygen through the electrolytic conversion of water and a cell-encapsulating pouch made of alginate-impregnated electrospun fibrous membrane for immunoprotection. Given biocompatibility is a pivotal requisite for the clinical translation of biomedical devices, all the components of the BEAM system were engineered from medical-grade materials. iEOGs were GMP-manufactured by Giner Inc. The electrolyzer components were housed within a medical-grade titanium enclosure with a porous polymeric membrane integrated onto one face, functioning as a window to harvest interstitial water vapor from surrounding tissue for electrolysis. The iEOGs were controlled to a settable current with corresponding voltages from 1.4 to 1.8 V, in the range for the splitting of water into hydrogen and oxygen, which were released through two separate outlets (**Figure 1A**). The oxygen outlet delivered gas by convection to an oxygen-permeable tubing traversing the cylindrical cell encapsulation pouch, enabling oxygen diffusion along its length to the encapsulated cells (**Figure 1B,C**). An alginate-impregnated fibrous membrane was used as the immuno-protective barrier, which allows for selective diffusion of nutrients, glucose, and insulin while preventing the infiltration of host cells.

**Figure 1.**
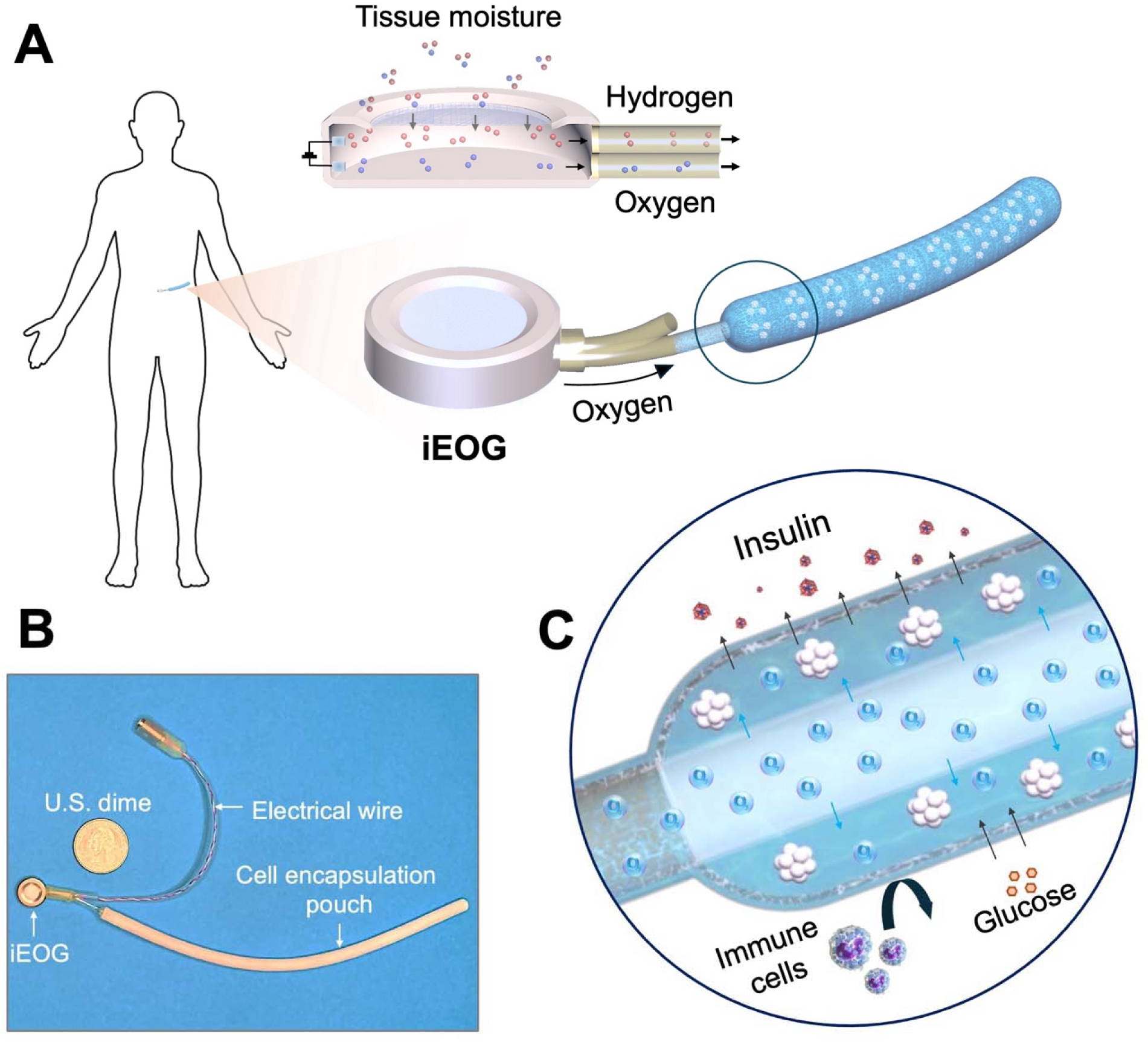
Conceptual design of the BEAM system. The system comprises two integral components: an implantable electrochemical oxygen generator (iEOG) and a core-shell cell encapsulation pouch designed in linear configuration to facilitate minimally invasive implantation, connected by gas-impermeable tubing. (**A**) Schematic representation illustrating oxygen generation by electrolytic splitting of tissue moisture. A human sketch is shown to indicate the relative size of a human version system. (The power supply needed for electrolysis is not shown) (**B**) A representative digital image of a scaled-up BEAM system with a cell encapsulation pouch measuring 15 cm in length and 5 mm in thickness. (**C**) Longitudinal section view of the cell encapsulation pouch showing pancreatic islets located in the annular space between an inner silicon tubing and the semipermeable membrane for optimal mass transportation and proximity between cells and the oxygen source.

Two generations of the iEOG were developed by Giner Inc based on the membrane electrode assembly (MEA) technology as previously described [29, 30] (**Figure 2A**). The MEA consists of a proton exchange membrane that when hydrated serves as the electrolyte, transferring the protons in the balanced electrochemical reaction. The anode and cathode electrocatalysts are pressed on opposite sides of the membrane as a single piece composite. At the surface of the anode, oxygen is evolved and leaves via convection through the oxygen gas port on the anode side. At the surface of the cathode, hydrogen is evolved and leaves via convection through the hydrogen gas port on the cathode side. The first version (v1 iEOG) had dimensions of 20 mm in diameter, 4.6 mm in height, and a weight of approximately 4 grams. The second version (v2 iEOG) was designed to be smaller in both diameter and thickness by decreasing the MEA area, while maintaining an equivalent oxygen generation capacity. Specifically, the v2 iEOG measures 13 mm in diameter, 3.1 mm in height, and approximately 2 grams in weight, which is approximately 25% of the volume and 50% of the weight of the v1 iEOG. Both versions demonstrated precise regulation of oxygen production via modulation of electrical current, allowing for variable oxygen generation rates tailored to meet the metabolic demands of different cells and cell doses, ranging from small numbers (∼1000 IEQ) to clinically relevant doses of islets. The average dose of cadaveric pancreatic islets transplanted into the portal vein during the U.S. CIT-07 trials (NCT00434811) was approximately 800,000 IEQ [31]. While the optimal dose of islets for extrahepatic transplantation sites has not yet been established, it was predicted that the required dose would range from 300,000 IEQ to 770,000 IEQ for a 70 kg patient [32]. To evaluate whether the oxygen generation capability of our system could meet the oxygen demand of clinically relevant doses of islets over extended periods, we operated three v1 iEOGs at 10 mA in saline for 11 months, 2 years, and 2.5 years. All three devices exhibited consistent currents close to the setpoint throughout the duration of the study, resulting in a stable oxygen output of approximately 1.94 scch (**Figure 2B**). This oxygen flow rate is sufficient to support >600,000 human IEQ, assuming an oxygen consumption rate of 2.1 pmol/min/IEQ (**Supplementary Table 1)**. To assess whether the iEOGs could achieve similar oxygen production *in vivo*, the devices were implanted in immunodeficient nude rats or immunocompetent Sprague-Dawley (SD) rats, operating at 11 mA to simulate expected clinical conditions. After three months of implantation in RNU nude rats, the v1 iEOG generated comparable oxygen levels (2.31 ± 0.10 scch) to those measured pre-implantation (2.25 ± 0.01 scch), which were calculated to be sufficient to support 720,000-740,000 human IEQ (**Figure 2C**). Similar results were observed for the v2 iEOG, with oxygen production levels of 2.29 ± 0.03 scch maintained after two months of implantation in nude rats (**Figure 2D**). In immunocompetent SD rats, the v2 iEOG also demonstrated stable oxygen generation, with levels measured at 2.44 ± 0.16 scch before implantation and 2.29 ± 0.12 scch one month post-implantation (**Figure 2E**). Taken together, these findings suggest the long-term capability of iEOGs to produce sufficient oxygen from interstitial moisture to support clinically relevant doses of human pancreatic islets. Furthermore, histological analysis using Masson’s trichrome staining of tissue surrounding the iEOG in SD rats after one month showed no evidence of adverse tissue reactions and indicated the presence of blood vessels near the water-harvesting window (**Figure 2F,G**). We selected the v2 iEOG for further development given the compact design and reduced weight, making it more suitable for implantation.

**Figure 2.**
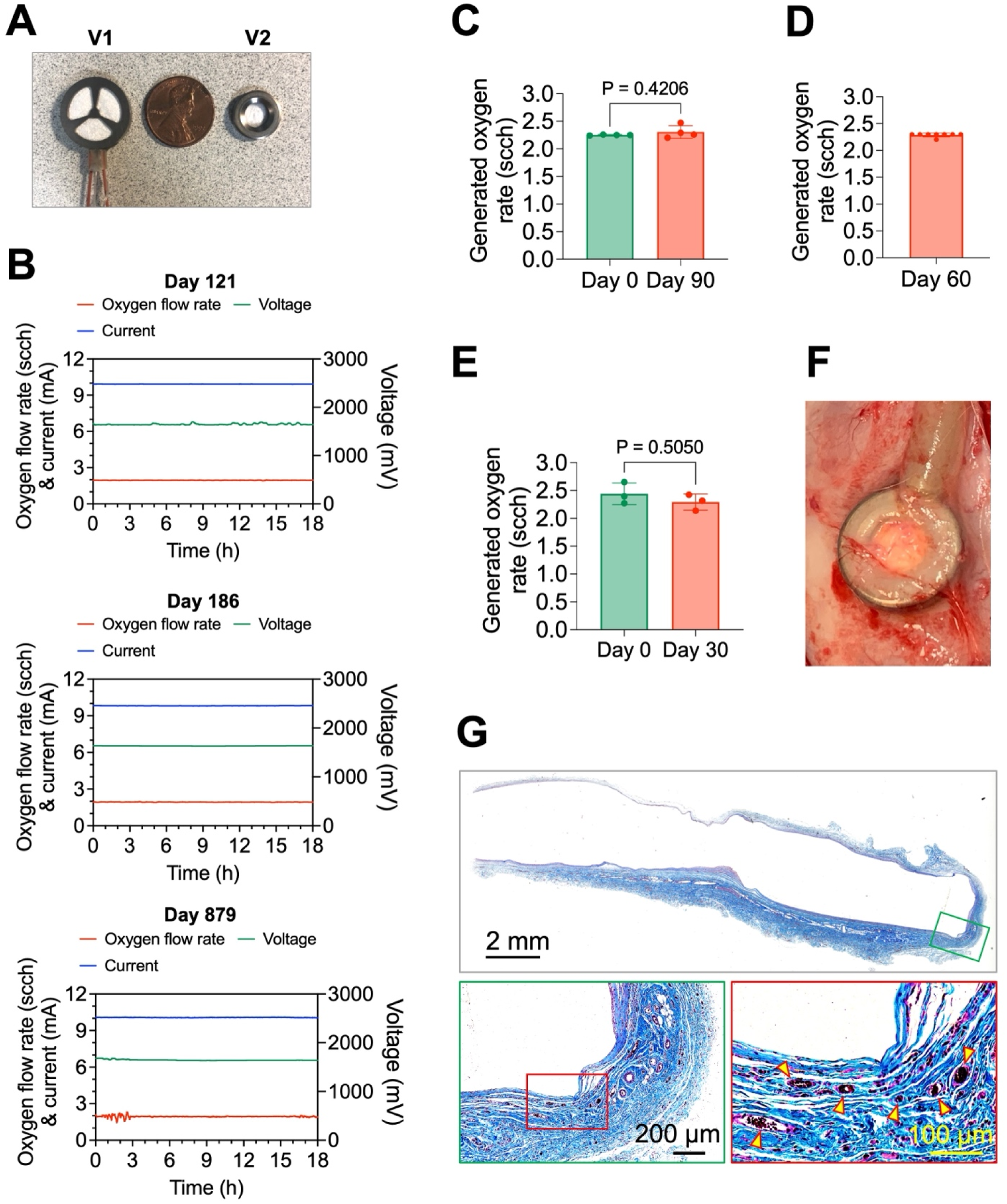
*In situ* oxygen generation and biocompatibility of iEOG. (**A**) A digital image comparing the dimensions of v1 iEOG and v2 iEOG. (**B**) Representative 18-hour measurements of oxygen flow rate, current, and voltage from a v1 iEOG operated at 10 mA over 2.5 years, collected on day 121, 186, and 879. (**C**) Oxygen generation rates of v1 iEOGs operated at 11 mA, measured before and 3 months post-implantation in RNU nude rats (n=4). Data are presented as mean ± SD. Statistical analysis was performed using paired two-tailed Student’s t-test. (**D**) Oxygen generation rates of v2 iEOGs operated at 11 mA, measured 2 months post-implantation in RNU nude rats (n=8). Data are presented as mean ± SD. (**E**) Oxygen generation rates of v2 iEOGs operated at 11 mA, measured before and 1 month after implantation in immunocompetent SD rats (n=3). Data are presented as mean ± SD. Statistical analysis was performed using paired two-tailed Student’s t-test. (**F**) A representative stereomicroscope image showing a v2 iEOG implanted in the subcutaneous space of SD rats after 1 month. The skin was removed to exposure the implant. (**G**) Masson’s trichrome staining of tissue surrounding the iEOG after 1 month of implantation in SD rats. Arrowheads indicate the presence of blood vessels.

The cell encapsulation pouch was designed to be mechanically robust, scalable, and biocompatible. Mechanical properties, specifically stiffness and elasticity, significantly impact the foreign body responses following implantation [33–36], which is a crucial determinant for the outcomes of a cell encapsulation implant. Ideally, the pouch should be soft to reduce the mechanical mismatch with the surrounding host tissue, which can trigger the activation of inflammatory and fibroblastic cells [37–39]. On the other hand, it also needs to be elastic and durable to maintain its structural integrity during surgical handling and long-term indwelling as intensified fibrotic responses were generally observed in the rough and kinked regions caused by implant deformation [40](**Supplementary Figure 2**). To meet these requirements, we used an electrospinning technique to prepare a soft, highly porous pouch with round end using medical grade polyether block amide (PEBAX) as a base material (**Figure 3A**). Arkema Pebax® thermoplastic elastomer was selected for its inherent softness (77 Shore D) and flexible, rubber-like nature (Flexural modulus of 12). Scanning electron microscopy (SEM) of the membrane revealed a fibrous structure with interconnected pores, which is essential for efficient mass transfer (**Figure 3B,C**). The fibers were randomly aligned with sizes ranging from 1.4 to 4 μm (average size: 2.47 ± 0.43 μm) (**Figure 3D**); we chose this relatively large fiber sizes to facilitate mass transfer and improve capillary-driven penetration of alginate into the membrane during the hydrogel coating. The porosity of the membrane was estimated to be 55.12 ± 5.93% (**Supplementary Figure 3**). The thickness of the membrane was approximately 70 μm, as determined by SEM. Furthermore, the membrane demonstrated considerable mechanical robustness and elasticity, as indicated by a tensile strength greater than 8 MPa and a fracture strain exceeding 23 (**Figure 3E**).

**Figure 3.**
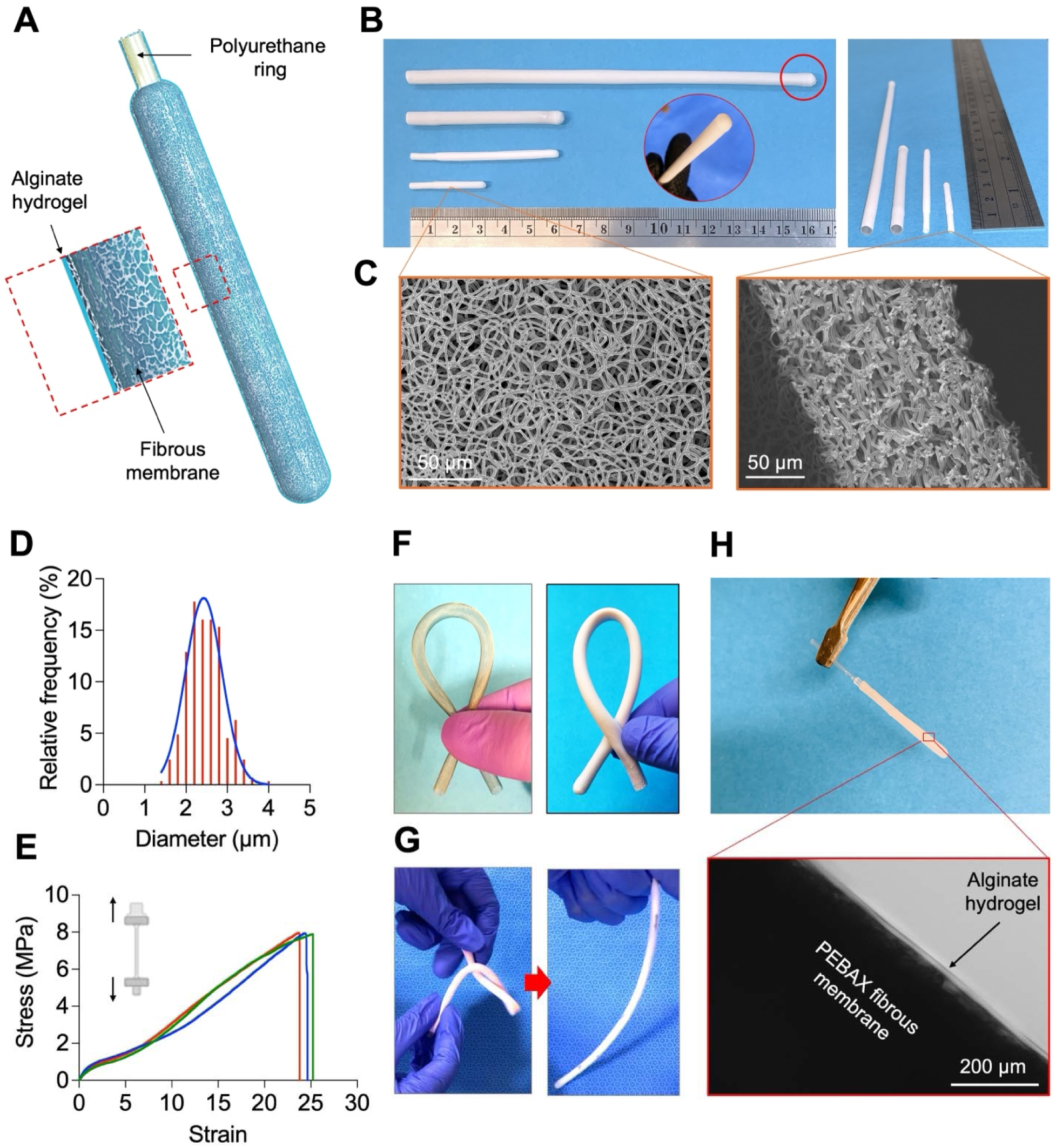
Characterization and scalability of the cell encapsulation pouch. (**A**) Schematic illustration for the structure of a cell encapsulation pouch. (**B**) Representative digital images showing cell encapsulation pouches in various lengths and diameters, designed to accommodate different cell doses. The inset shows the round end of the pouch. (**C**) SEM images of the pouch surface and cross-section. (**D**) Size distribution of PEBAX fibers. (**E**) Tensile testing of PEBAX cell encapsulation pouches (n=3). (**F**) Digital images of a nitinol scaffold and a nitinol-reinforced pouch. Both were bent without kink. (**G**) Digital images demonstrating the ability of nitinol-reinforced pouches to reverse to their original structural form following twisting deformation. (**H**) Representative microscopy image showing the alginate impregnation onto the encapsulation pouch.

One of the challenges in the fabrication of membrane-based encapsulation systems is achieving effective sealing. Conventional sealing methods involving the use of heat, ultrasonic welding, or adhesives can be troublesome due to interference from liquid (media, crosslinking solution) used during the cell loading process. In addition, these methods generally result in rigid and sharp edges, which potentially exacerbate the fibrotic responses. The by-products generated during the application of heat or glue may further contribute to these adverse reactions. Indeed, significantly enhanced fibrotic responses were observed at the heat-sealed ends of membrane-based systems compared to the membranes themselves, regardless of the membranes’ physicochemical properties, across various immunocompetent animal models, including C57BL/6 mice (**Supplementary figure 4-7**), SD rats, and Göttingen minipigs [41]. The necessity of employing heat sealers, welders, or adhesives also complicates the fabrication process, especially under aseptic conditions. To address these challenges, we designed an edge-free cell pouch with a specific geometry to minimize the area requiring sealing (**Figure 3A**). A water-soluble polyethylene glycol (PEG) template was employed to collect the electrospun fibers, thus shaping the cell encapsulation compartment into an edge-free round-ended pouch (**Supplementary Figure 8**). A medical-grade polyurethane ring was incorporated into the other open end to serve as a female component in a barbed slide-in connection. This configuration allows the pouch to be sealed by simply sliding the oxygen transportation tubing into the lumen of the cell encapsulation pouch.

Figure 3B shows that the encapsulation pouch can be scaled both radially and longitudinally to accommodate different doses of cells. For long pouches (e.g. human version) which tend to kink, the membrane was reinforced with an elastic, shape-memorable nitinol braided mesh scaffold with a wire thickness of 25.4 μm and pore sizes ranging from 100 μm to 150 μm (Figure 3F; **Supplementary Figure 9; Supplementary Video 2**). The reinforced pouches did not kink and were able to revert to their original form after common deformations, such as folding, bending, or twisting, that may occur during the surgical procedure or host movement (Figure 3G**; Supplementary Video 3**). The impact of the nitinol scaffold on mass transfer is expected to be negligible owning to the large pore sizes and thin wire thickness. To impart the immuno-protective property, the fibrous membrane was impregnated with clinical-grade alginate (SLG100) using a previously developed method to avoid the detachment of alginate hydrogel after implantation [42]. This resulted in a hydrogel layer thickness of 30-50 μm, bringing the final thickness of the pouch wall to approximately 100-120 μm (Figure 3H).

### Cell Loading and System Assembly

A customized cell-loading apparatus, constructed from a perforated polytetrafluoroethylene (PTFE) tube, was developed for cell loading into the BEAM system. PTFE was chosen due to its inertness and hydrophobic properties, which effectively reduce the adhesion of residual alginate solution, thereby minimizing cell loss during encapsulation. To demonstrate the loading procedure, we used INS-1 cell aggregates with a size distribution from 50 to 300 μm as a model to mimic pancreatic islets (**Supplementary Figure 10**). These aggregates were gently dispersed in a 2% alginate solution (SLG100) at a density of 60,000 IEQ/mL before being introduced into the cell-loading apparatus. **Figure 4A** shows the streamlined procedure of cell loading and system sealing. To better visualize the cell loading process, we loaded 150-μm beads (used to simulate cell aggregates) through the loading apparatus, into an encapsulation pouch made of a transparent tube, resulting in a relatively even distribution within the annular space between the loading tool and the pouch wall (**Supplementary Figure 11**). When the encapsulation pouch was immersed in a crosslinking solution containing 95 mM calcium chloride and 5 mM barium chloride, the divalent ions diffused through the porous membrane, initiating alginate gelation. Both longitudinal and cross-sectional histology images of the pouches revealed the annular configuration of the aggregate-laden alginate layer with the thickness of 300-500 μm, which is equivalent to about 2-3 layers of cell aggregates (**Figure 4B**). This thin hydrogel layer was found to adhere to the fibrous membrane, thus reducing the distance between the cells and the pouch surface to enhance mass transfer. Live/dead staining and histological examination showed no significant changes in aggregate morphology or viability after loading into the BEAM system (**Figure 4C**).

**Figure 4.**
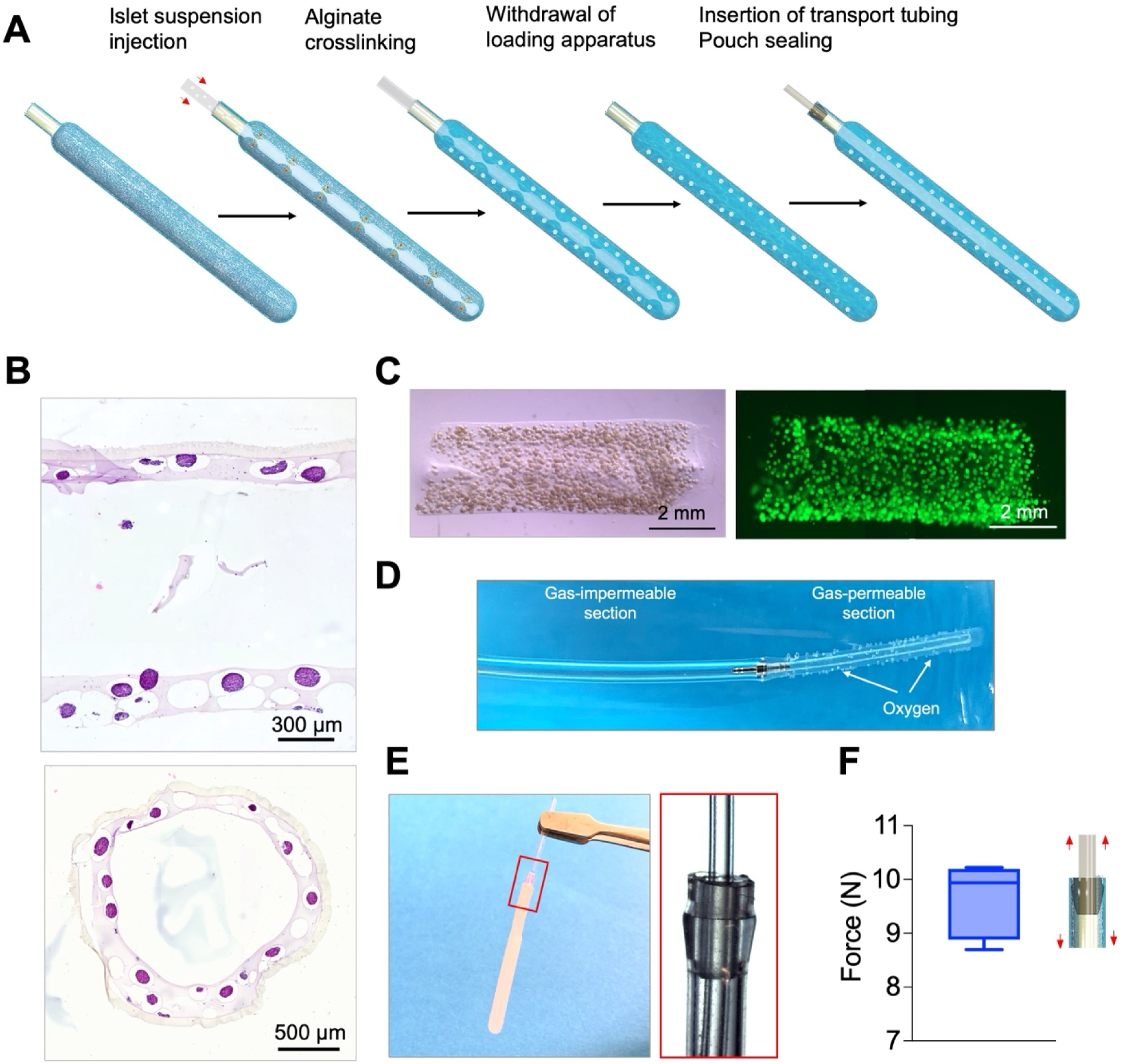
Cell loading and encapsulation pouch sealing. (**A**) A schematic diagram illustrating stepwise procedure for cell loading and pouch sealing in the BEAM system. (**B**) Hematoxylin and eosin (H&E) staining of longitudinal and cross-sections of an encapsulation pouch loaded with INS-1 cell aggregates at 60,000 IEQ/mL. (**C**) Bright-field and fluorescence images of the hydrogel layer removed from the encapsulation pouch, containing green fluorescence-labeled INS-1 cell aggregates at 60,000 IEQ/mL. Scale bar: 2 mm. (**D**) A representative digital image of an oxygen transportation tubing infused with a high flow rate of oxygen. (**E**) Representative images depicting the connection between the oxygen transportation tubing and the cell encapsulation pouch. The fibrous membrane was removed for visualization purpose. (**F**) The pull-off force measured during the tensile test, indicating the force at which the barb connector is removed from the cell encapsulation pouch (n=5). The data are presented as a min to max box-and-whisker plot.

Upon crosslinking of the alginate layer, the loading apparatus was withdrawn, allowing for the insertion of the oxygen-transportation tubing. This tubing comprised end-sealed silicone tubing connected to gas-impermeable polyurethane tubing, thereby facilitating oxygen diffusion selectively through the silicone wall to support adjacent cells (**Figure 4D**). A barbed connector was integrated onto the gas-impermeable section, designed to fit with the polyurethane ring of the cell encapsulation pouch. Once the oxygen-transportation tubing was fully inserted, the barbed connector provided a tight fit with the ring, effectively sealing the pouch to prevent gas leakage and maintain immunoprotection (**Figure 4E**). The connection demonstrated substantial robustness, with the ability to withstand pull forces exceeding 8.5 N (**Figure 4F**). The sealed cell encapsulation pouch was finally connected to an iEOG using a custom barb connector before implantation.

### Oxygenation by the BEAM system preserved the viability and function of INS-1 aggregates under an *in vitro* hypoxic condition

The local partial oxygen tension surrounding an encapsulation implant is a critical determinant for the survival and function of encapsulated β-cells, particularly when these cells are loaded at a high density. Unfortunately, post-transplantation oxygen levels in the vicinity of subcutaneous implants have been reported to be critically low (<10 mmHg), primarily due to inadequate vascularization at the subcutaneous implantation site and the subsequent formation of a fibrotic capsule [43–45]. Given the inherently high oxygen consumption rate of β-cells, such low oxygen tension is insufficient to support their long-term survival and function. In addition, this also limits the feasibility of loading them at a high density, thus rendering the requisite implant size impractical for clinical applications. We postulated that enhancing the oxygen supply to the implant could improve cell survival and elevate the upper limit of islet cell density. To test this hypothesis, we encapsulated INS-1 832/13 aggregates (2000 IEQ), an insulin-secreting cell line with oxygen consumption rates comparable to that of rat pancreatic islets and higher than that of human islets (**Supplementary Table 1**), in a BEAM system, resulting in a relatively high loading density (60,000 IEQ/mL). These systems were challenged under 1% O_2_ cell culture condition (∼ 7.62 mmHg) for 24 hours (**Figure 5A,B**). We observed significant cell death in non-oxygenated implants, as evidenced by cell disintegration, shrunken morphology, and live/dead staining (**Figure 5C,D**). Histological analysis further revealed a fragmented aggregate structure and cell apoptosis evidenced by weak cytoplasmic staining, nuclear shrinkage, and the formation of apoptotic bodies in the majority of cells (**Figure 5E**). Additionally, insulin-positive cells were scarcely detected in these non-oxygenated systems (**Figure 5F**). These findings demonstrate a substantial reduction in the viability and function of INS-1 832/13 cells following 24 hours of culture under a hypoxic condition. In contrast, cell aggregates in systems supplied with an oxygen generated at an electrical current of 270 μA maintained normal morphology and healthy aggregate structure, with a negligible dead cell number, as indicated by both live/dead staining and histological examination (**Figure 5B-F**). The oxygen generation rate was approximately 0.016 scch, as estimated using our simulation. In addition, the majority of cells in this group were positive for insulin staining, suggesting that oxygenation by the BEAM system effectively preserved INS-1 832/13 cell aggregate function under a hypoxic culture condition, even at a relatively high cell density.

**Figure 5.**
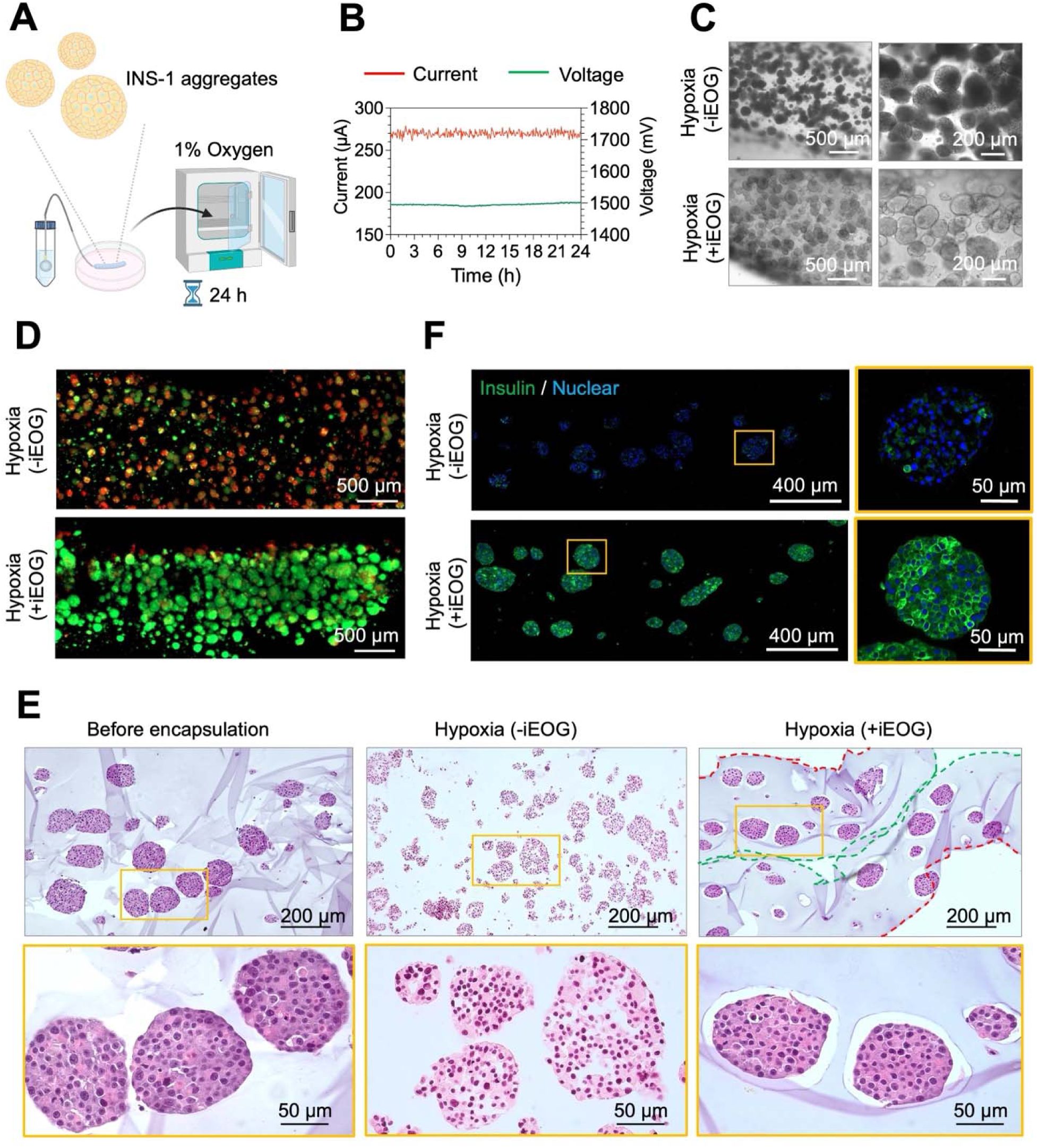
The impacts of oxygenation on INS-1 cell aggregates encapsulated in the BEAM system under a hypoxic cell culture condition. **(A)** A schematic representation of the experimental setup. (**B**) Operating electrical current and voltage of iEOG recorded during a 24-hour experiment. (**C**) Bright-field microscopic images of INS-1 cell aggregates in BEAM systems following a 24-hour incubation period under a hypoxic culture condition (1% oxygen), with and without supplemental oxygenation provided by iEOG. **(D)** Dual fluorescence staining of INS-1 cell aggregates within BEAM systems following 24-hour incubation in 1% oxygen, with and without additional oxygen supply from iEOG. Green: Acridine Orange, Live cells; Red: Ethidium homodimer, Dead cells. (**E**) H&E staining of INS-1 cell aggregates before encapsulation and those loaded in BEAM systems subjected to a hypoxic culture condition (1% oxygen) for 24 hours, with or without oxygen supplementation from iEOG. Green dashed line represents the interface with the silicone tubing while the red dash line indicates the interface with the fibrous membrane. (**F**) Immunofluorescence staining of INS-1 cell aggregates in BEAM systems following 24-hour incubation in 1% oxygen, with and without additional oxygen supply from iEOG. Green: Insulin; Blue: Nuclear staining. **Figure 5A** was created using BioRender.

### Oxygenation by the BEAM system preserved the viability and function of human islets under an *in vitro* hypoxic condition

We then assessed the impact of oxygenation provided by the BEAM system on the viability and function of primary human pancreatic islets. Approximately 1,800 human IEQ were dispersed in 30 μL of 2% SLG100 solution and subsequently loaded into a BEAM system at a density of 60,000 IEQ/mL and subjected to 1% O_2_ culture condition (**Figure 6A,B**). The BEAM system was operated at 240 μA, corresponding to an oxygen generation rate of approximately 0.009 scch, as estimated by our simulation. After 24 hours, pancreatic islets encapsulated in systems without oxygenation exhibited significant cell dissociation and apoptosis (**Figure 6C,D**). Immunofluorescence staining revealed a notable decline in cell functionality, as indicated by dominant loss in insulin and glucagon expression compared to those of healthy islets (**Figure 6E**). The impaired function was further corroborated by glucose stimulated insulin section (GSIS) assays (**Figure 6F**). The levels of insulin secreted from systems cultured under the normoxic condition for 24 h measured 405.6 ± 16.6 μIU/h and 745.9 ± 228.0 μIU/h in the 2.8 mM and 28 mM glucose solutions, respectively. Meanwhile, non-oxygenated systems exhibited significantly lower insulin secretion levels, with 25.5 ± 4.8 μIU/h in a 2.8 mM glucose solution and 99.1 ± 80.9 μIU/h in a 28 mM glucose solution. In contrast, live/dead staining and histological analysis demonstrated that pancreatic islets encapsulated in systems supplied with oxygen remained viable after 72 hours of incubation under hypoxic culture condition (**Figure 6C,D**). GSIS and histological evaluations indicated that the function of these islets was comparable to those incubated under normal culture conditions. After 24 hours of hypoxic incubation, insulin secretion levels from these oxygenated systems were 330.7 ± 92.0 μIU/h in the 2.8 mM glucose solution and 1,025.5 ± 172.2 μIU/h in the 28 mM glucose solution. Notably, strong insulin and glucagon staining were observed in pancreatic islets of this group after 24 h of incubation (**Figure 6E**). These findings suggest that the continuous supply of oxygen by the BEAM system effectively preserved the viability and function of human pancreatic islets encapsulated at a density of 60,000 IEQ/mL.

**Figure 6.**
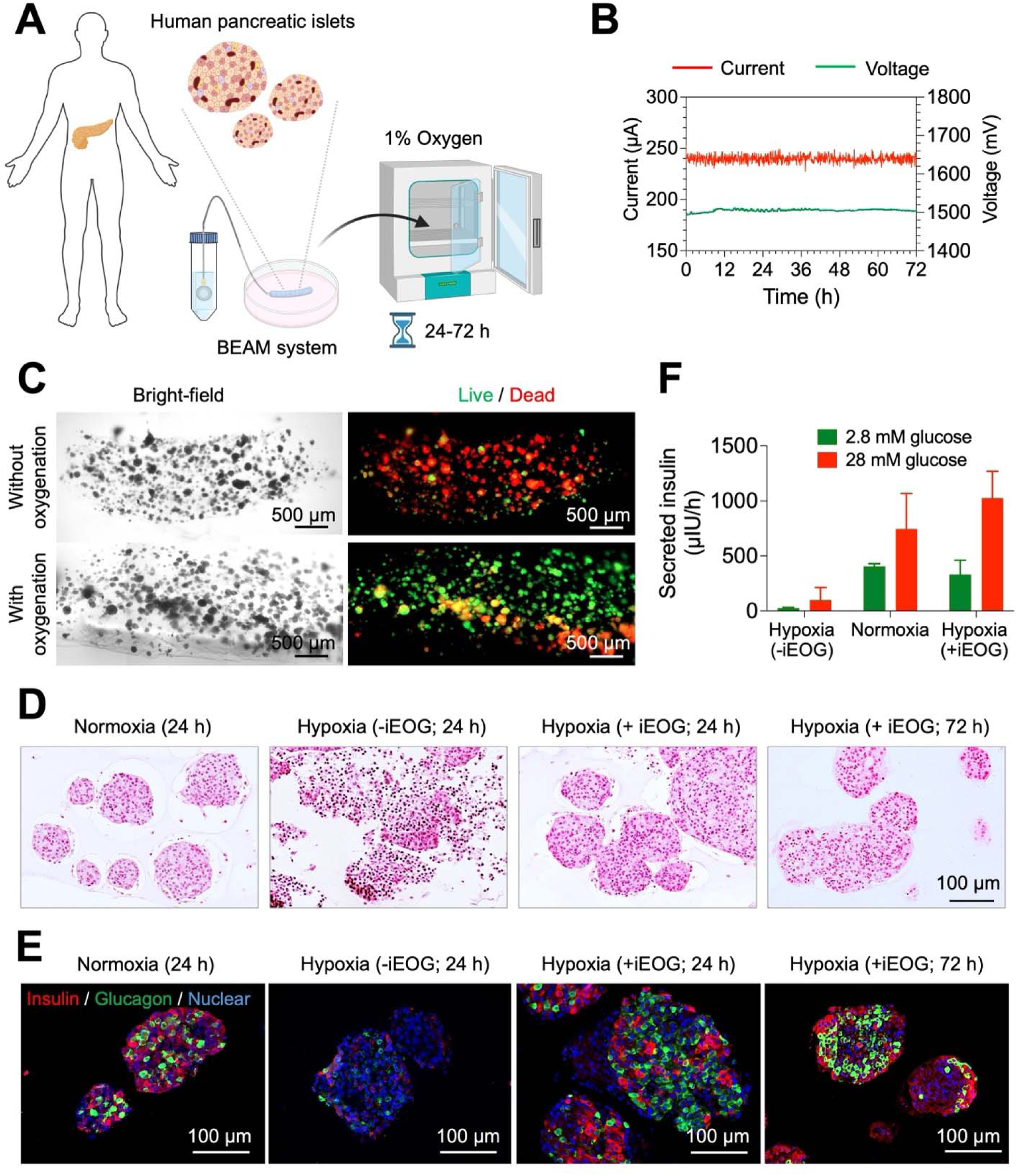
The impact of oxygenation on viability and function of human pancreatic islets encapsulated in BEAM systems under a hypoxic cell culture condition. **(A)** A schematic representation of the experimental setup. (**B**) Operating electrical current and voltage of iEOG recorded during a 72-hour experiment. (**C**) Dual fluorescence staining of islets encapsulated in BEAM systems subjected to a 24-hour incubation under hypoxic condition (1% oxygen), with and without supplementary oxygenation from iEOGs. Scale bar: 500 μm. Green: Acridine Orange, Live cells; Red: Ethidium homodimer, Dead cells. Dead cells in bottom right panel are thought to be a cutting artifact from processing the annual hydrogel. (**D**) H&E staining of encapsulated islets after incubation under a normoxic condition and a hypoxic condition, with and without supplementary oxygenation from iEOG. (**E**) Immunofluorescence staining of the encapsulated islets after 24-hour incubation under a normoxic and a hypoxic culture condition, with and without supplementary oxygenation from iEOG. Red: Insulin; Green: Glucagon; Blue: Nuclear staining. (**F**) Quantification of insulin secretion from islets encapsulated in BEAM systems, following a 24-hour incubation under normoxic condition (20% oxygen) (n=2) and hypoxic condition (1% oxygen), with and without oxygen supplementation from the iEOG (n=2). **Figure 6A** was created using BioRender.

### Effects of oxygenation by the BEAM system in an allogeneic rat transplantation model

After validating the favorable effect of oxygenation *in vitro*, we extended our investigation to assess its potential to enhance the viability and function of pancreatic islets *in vivo* using an allogeneic rat model. Pancreatic islets (6000 IEQ) isolated from Lewis rats were encapsulated in a cell pouch (3.2 mm O.D. x 2 cm length; loading capacity of ∼100 μL) of the BEAM system and transplanted into STZ-induced diabetic, immunocompetent SD rats. All rats selected for implantation exhibited blood glucose levels exceeding 450 mg/dL for over 7 consecutive days. The weights of rats before implantation were 418.8 ± 58.5 grams, resulting in the islet doses of 14,509.5 ± 1,729.6 IEQ/kg. Owing to the relatively large size of the current electronic controller (3.5 cm x 3.5 cm x 1.2 cm) designed for bench testing, it was not feasible to fully implant the entire BEAM system within the subcutaneous space of rats. Thus, we used an extracorporeal setting, in which electronic components including battery, electronic controller, and iEOG enclosed in a refillable water reservoir, were secured in a pocket worn by the rats (**Figure 7A,B; Supplementary Figure 13**). This setting enables convenient monitoring of iEOG function, water consumption, and battery exchange. For instance, the extracorporeal electronic box allows for the incorporation of a light-emitting diode (LED) to indicate the status of oxygen production for early detection of electrolyzer malfunction or system disconnections, which is crucial in long-term animal study. It is worth noting that for clinical applications, the entire system would be implanted with rechargeable power supply charged weekly via transcutaneous energy transfer (see **Supplementary Figure 14**). For the rat study, we designed a customized animal jacket with a small pocket on the back to secure these electronic components. The flank and abdominal area of the jacket were left open to minimize the discomfort or stress on the rats (**Supplementary Figure 15**). The use of jackets did not significantly impair the mobility of rats, nor did it cause negative health issues, as evidenced by steady body weight gain over a month (**Supplementary Figure 16**). The battery was exchanged every 5 days, and the water reservoir was replenished on a weekly basis. The iEOGs were operated at 280 μA to achieve oxygen generation rates of 0.455 ± 0.015 scc per day (**Figure 7 C, D**). The cell encapsulation pouch was implanted subcutaneously through a minor skin incision and connected to the iEOG via percutaneous oxygen-impermeable polyurethane tubing. A medical-grade polyester tissue cuff was incorporated at the incision to aid healing and prevent infection. In all 9 animals that underwent transcutaneous implantation, we observed complete wound closure without notable signs of inflammation or adverse reactions at the incision site (**Supplementary Figure 17**). Implants without connection to an iEOG (4/4) failed to reverse diabetes, whereas all rats (9/9) received an iEOG-connected implant returned to normoglycemia within 3 days (**Figure 7E-G**). Early oxygenation cessation occurred in 3 rats from the first cohort on day 1, day 3, and day 5 because of the damage of the transcutaneous tubing and electronic wire, which were exposed out of the jackets and compromised by the animals (**Supplementary Figure 18; Supplementary Table 2**). The implantation location and jacket were then adjusted so these components were secured under the jacket. As a result, the remaining rats with continuous oxygenation maintained normoglycemia for over a month and up to 88 days. Intraperitoneal glucose tolerance test (IPGTT) on day 35 indicated that these rats tolerated glucose comparably to healthy rats, suggesting the normal metabolic function of encapsulated islets (**Figure 7H,I; Supplementary Figure 19**) without significant mass transport limitations. This was further corroborated by the significant increases in serum C-peptide after glucose administration (**Supplementary Figure 20**) and the body weight differences between the oxygenated and non-oxygenated groups (**Figure 7G**). In contrast, the blood glucose levels of rats implanted with non-oxygenated implants remained higher than 400 mg/dL after 120 min. Oxygenation was accidentally stopped in 2 rats on day 51 and day 72 due to the disconnection of the iEOG with the electronic controller and the cell pouch caused by the rats (**Supplementary Figure 18; Supplementary Table 2**). Intriguingly, sharp increases in blood glucose levels were observed right after oxygenation cessation at both early time points (day 1, 3, and 5) and later time points (day 51 and day 72). This suggests that a continuous supply of oxygen may be vital for the long-term function of pancreatic islets in cell encapsulation systems, particularly at high loading density. This is in contrast to a hypothesis in the encapsulation field that short-term oxygenation in the early period may be sufficient.

**Figure 7.**
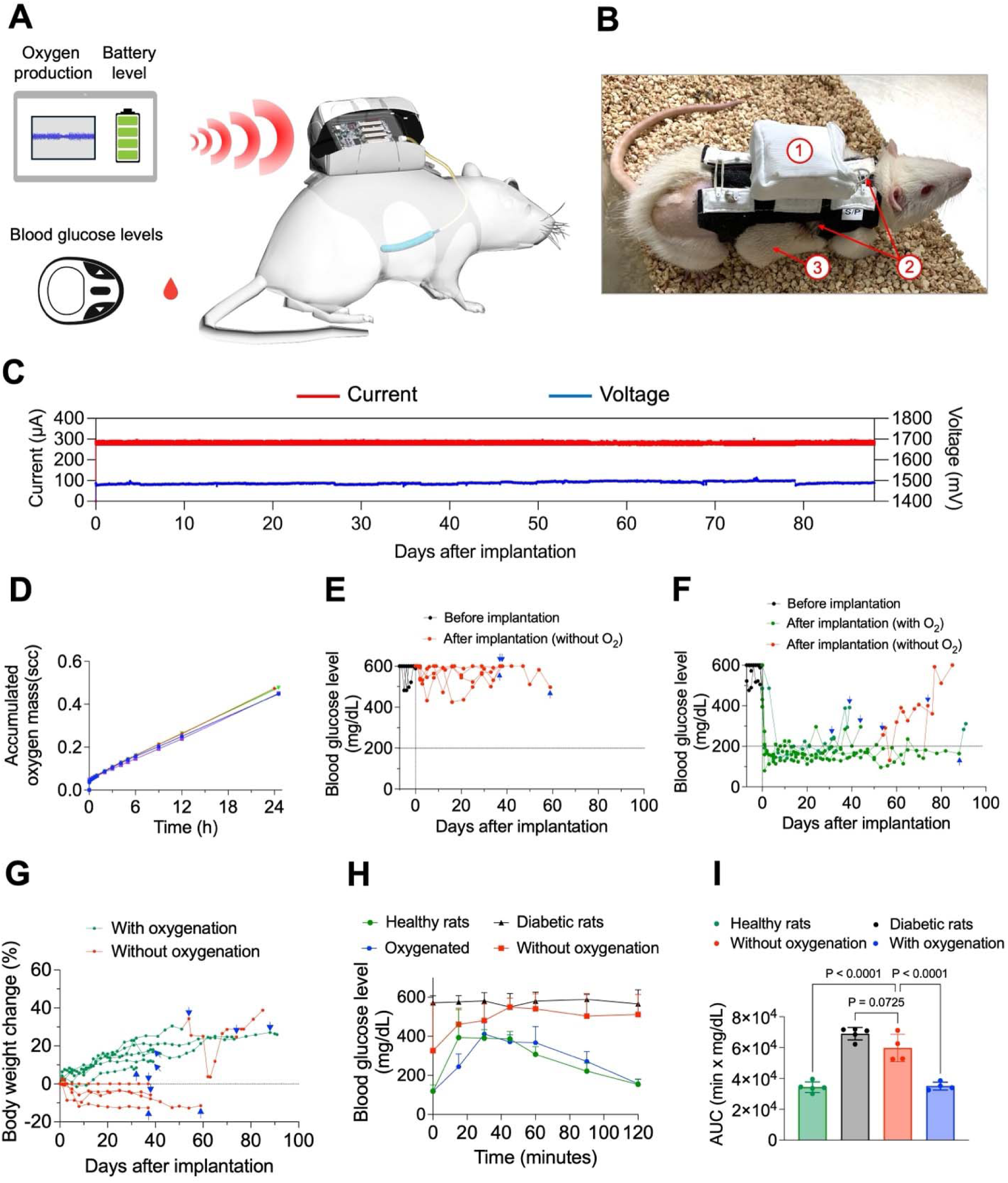
The impact of oxygenation on the metabolic function of pancreatic islets encapsulated within the BEAM system in an allogeneic rat model. (**A**) A schematic illustration of experimental design. Current, voltage and battery level were wirelessly spot measured via IR communication. (**B**) A digital image demonstrating the BEAM system assembly on a diabetic rat. (1) A pocket on the back of a jacket worn by the rat to secure the iEOG and an electronic box. (2) The oxygen transportation tubing running from iEOG, underneath the jacket to the (3) cell encapsulation pouch implanted in the subcutaneous space of the rat. (**C**) Operating current (red) and voltage (blue) of the iEOG recorded during 88-day study from the electronics memory. (**D**) Cumulative volume of oxygen generated from the iEOGs used on the rats over a 24-hour period (n=4). (**E**) Non-fasting blood glucose levels of rats implanted with a cell encapsulation pouch without oxygen supplementation (n=4). (**F**) Non-fasting blood glucose levels of rats received implantation of a BEAM system (n=6). (**G**) Body weight changes of rats received implants with or without supplemental oxygenation. In F and G, green dots represent blood glucose levels/body weight changes during continuous oxygenation, while red dots represent blood glucose levels/body weight changes measured without oxygenation or after the oxygenation cessation. Blue arrows mark the timepoint of implant retrieval. (**H**) Intraperitoneal glucose tolerance test conducted on healthy rats (n=5), diabetic rats (n=5), rats receiving a BEAM system (n=4; measured on day 35), and rats receiving a cell encapsulation pouch without oxygen supplementary (n=4; measured on day 35). Data are presented as mean ± SD. (**I**) Corresponding area under the curve for the different groups. Data are presented as mean ± SD. Statistical analyses were performed using one-way ANOVA.

Post-retrieval evaluations further revealed prominent cell death with shrunken morphology in implants without oxygenation and implants with ceased oxygenation, as indicated by both live/dead staining and histology examination (**Figure 8A-D; Supplementary Figures 21-22**). Almost no insulin-positive cells were found in these implants (**Figure 8C**). No signs of host cell infiltration were observed in these systems, suggesting the destruction of pancreatic islets was not likely caused by immune cell-mediated rejection. GSIS assay demonstrated low and no significant differences between insulin levels secreted by these cells under a low glucose condition (2.8 mM) and a high glucose condition (16.7 mM), suggesting compromised function of β-cells (**Figure 8E**). In contrast, viable islets with healthy morphology and strong insulin/glucagon staining were found in the oxygenated implant on day 32, 40, 45, and 88, suggesting the feasibility of oxygenation to preserve viability and function. GSIS assay on the islets retrieved from these implants showed substantial levels of secreted insulin and the ability to respond to glucose level change. It is worth noting that among these implants, three were retrieved after the blood glucose levels started to increase. We observed a gas bubble partially surrounding the implants in the rats with escalated blood glucose levels (3/6), probably due to the accumulation of excessive oxygen that separated the implants from the surrounding tissue (**Supplementary Figure 23**). To confirm this hypothesis, excessive gas was vented through a 23G needle puncture in a rat, quickly restoring normal blood glucose levels within a few hours and maintaining normoglycemia for an additional 2 weeks until oxygenation accidentally stopped on day 72.

**Figure 8.**
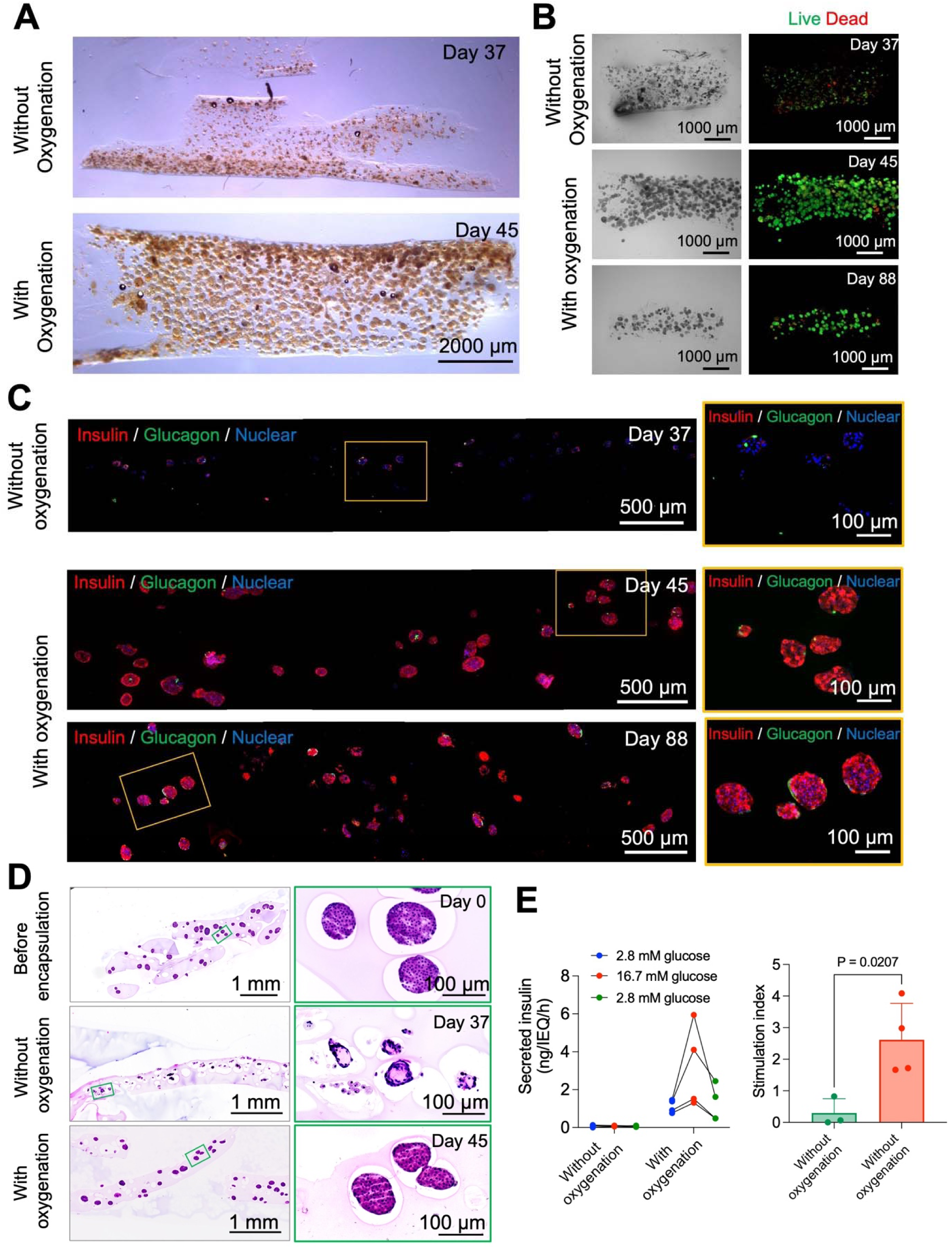
Oxygenation by the BEAM system effectively preserved the viability and function of encapsulated pancreatic islets in an allogeneic rat model. (A) Stereomicroscopic images showing islet-laden alginate hydrogel layer peeled off from an implant without oxygenation and an implant with oxygenation. (B) Viability assessment of encapsulated islets by dual fluorescent staining. Green: Acridine Orange, Live cells; Red: Ethidium homodimer, dead cells; Blue: Nuclear staining. (C) Representative immunofluorescence staining of encapsulated islets in implants without oxygen supplementation and implants with oxygen supplementation. Red: Insulin; Green: Glucagon; Blue: Nuclear staining. (D) H&E staining of islets before encapsulation, islets in an implant without oxygenation, and islets in an implant with oxygenation. (E) *Ex vivo* GSIS of encapsulated islets in implants without oxygen supplementation (n=3; day 37, 38 and 59) and implants with oxygen supplementation (n=4; day 32, 40, 45, and 88). Data are presented as individual values (left) and mean ± SD (right).

## Discussion

Macroencapsulation remains a compelling strategy for cell therapy in the treatment of T1D as the use of a permselective membrane could potentially obviate the need for chronic immunosuppression. An increasing body of successful proof-of-concept preclinical studies in mice has demonstrated the potential of this approach to protect encapsulated insulin-secreting cells from immune rejection, thereby enabling long-term glucose homeostasis without the need for immunosuppression [46–54]. However, the translation of these findings to larger animal models, as well as the scaling up of implants for clinical applications, remains a significant challenge. While a few hundred pancreatic islets, loaded within reasonably sized implants at low densities, may be sufficient to reverse diabetes and maintain normoglycemia in mouse models, scaling up these implants for use in larger animals and eventually humans is considerably more complex. The complexity arises from the necessity to pack islets at a high density to achieve an implant size that is practical for implantation. Pancreatic islets have a well-documented reliance on oxygen for their insulin secretion in response to glucose changes. Beta cells can survive under partial oxygen tension levels exceeding 0.1-0.44 mmHg while levels greater than 10 mmHg are necessary to sustain their ability for insulin secretion [8, 55]. Unfortunately, the local oxygen tension within encapsulation systems is significantly below the threshold required for the full function of pancreatic islets when transplanted into the poorly vascularized subcutaneous space [43–45]. This is further exacerbated by the additional mass transport barrier caused by the membrane and the subsequent formation of a fibrotic capsule, which is known to be more severe in larger animals and humans. Results from clinical trials by Viacyte Inc with PEC-Encap™ and PEC-Direct™ suggest that even with the use of membranes designed to promote vascularization or the membrane with perforations, the devices may fail to achieve sufficient vascularization in both timing and density to support a metabolically adequate functional beta cell mass [56]. Moreover, increasing cell density further heightens oxygen demand due to the steeper oxygen gradients within cell layers. Thus, in the previously developed cell encapsulation systems, insulin-secreting cells were typically loaded at densities not exceeding 25,000 IEQ/mL (**Supplementary Table 3**). This would necessitate impractically large encapsulation systems to accommodate a clinically relevant dose of pancreatic islets (300,000 IEQ-770,000 IEQ) at such low densities. For instance, in the clinical trial by ViaCyte (NCT04678557), 12 planar devices with a surface area of approximately 10 cm^2^ for each were required to achieve a metabolically relevant dose of cells. Computational modeling simulations suggest that a fiber measuring 12.8 meters in length with a 1 mm diameter, or a circular slab with a diameter of 11.9 cm and a thickness of 0.5 mm, would be required to deliver 500,000 functional IEQ under 40 mmHg O [57]. Even with microencapsulation, which provides a higher surface-to-volume ratio as compared to macroencapsulation systems, rapid decline in functionality was observed when the loading density was increased to 16,000 IEQ/mL [54]. Thus, oxygen supplementation is likely needed to realize the clinical translation of cell encapsulation.

Recent advancements in cell encapsulation technology have increasingly focused on the development of encapsulation systems capable of providing oxygenation to preserve viability and function of encapsulated cells [16, 18, 25, 58, 59]. For examples, metal peroxides [16, 18, 22, 24, 25] have been employed to generate oxygen to enhance cell survival in several encapsulation platforms. However, the inherently short-term oxygen supply in these methods poses a significant barrier to clinical translation, which demands long-term oxygenation. Thus, the need for a method to provide a continuous and sustainable oxygen source remains critical. A recent proof-of-concept study by Krishnan et al. successfully demonstrated the feasibility of using electrolysis-based oxygenation to improve the survival of encapsulated cells in subcutaneous space [59]. Their device, powered and regulated by resonant inductive coupling, effectively supported a moderate density of pancreatic islets (1000 IEQ/cm^2^) to correct diabetes in diabetic mice for a month. However, the practicality of resonant inductive coupling in clinical settings for generating an adequate level of oxygen to supply a metabolically relevant dose of cells requires further development and investigation.

In this study, we developed a bioelectronics-assisted encapsulation (BEAM) system incorporating a built-in battery and electronic controller, capable of generating oxygen in a continuously regulated and precise manner. The BEAM system is designed to overcome the limitations of oxygen supply duration associated with chemical-based methods by providing a theoretically unlimited oxygen supply through the electrolysis of tissue moisture, powered by a rechargeable energy source. The oxygen generator, with compact dimensions (13 mm diameter x 3.1 mm thickness) at its full scale, was proven to provide a sustained and adequate oxygen supply to support the upper limits of clinically relevant islet doses (600,000 IEQ-740,000 IEQ) for 11 months to 2.5 years *in vitro* and three months *in vivo*. The battery could be designed to be recharged weekly in a wireless fashion via transcutaneous energy transfer (TET), an established technology already in use for FDA cleared rechargeable neurostimulators, thereby reducing the need for patient intervention and compliance requirements. Furthermore, the oxygen generation rate is programmable and precisely controlled, allowing for adaptation to various cell doses and types with differing oxygen consumption rates.

Membrane electrode assembly (MEA) technology was used for the electrolysis of water to facilitate oxygen generation in this system. The MEA comprises a proton exchange membrane that, upon hydration, functions as the electrolyte, enabling proton transfer in the balanced electrochemical reaction. Hydrogen and oxygen evolve on opposite sides of the membrane and are transported through separate gas ports via convection, with less than 1% diffusive crossover, thereby minimizing the risk of spark ignition within the gas phases. The exterior of the iEOG is electrically insulated from the internal components, ensuring the absence of current or voltage hazards upon direct contact. Furthermore, the electrocatalyst-containing electrodes remain isolated from body fluids, as they receive water exclusively in vapor form through a vapor transport membrane, thereby mitigating any potential safety risks.

The linear outline of the BEAM system facilitates minimally invasive implantation and retrieval through a small skin incision. The cell encapsulation compartment is designed as an edge-free, flexible, round-ended cylindrical pouch, which minimizes the risk of enhanced fibrotic reactions typically caused by rigid, sharp edges. A straightforward sequential method for cell loading and system sealing was also developed, streamlining the preparation process, particularly under aseptic conditions. The concentric design of the cell encapsulation pouch, wherein cells are loaded into the thin annular space between the immune-protective membrane and the oxygen source (oxygen-permeable tubing), promotes optimal mass transport and ensures rapid, uniform oxygen diffusion. In addition, this structural configuration is scalable both radially and longitudinally, allowing for loading high islet doses while maintaining proximity between the encapsulated cells and the implant surface as well as the oxygen source.

Our initial results demonstrated that oxygenation provided by the BEAM system successfully preserved the viability and insulin secretion of INS-1 aggregates and human pancreatic islets under a severe hypoxic *in vitro* condition (1% oxygen or 7.65 mmHg). We then selected rats as the animal model due to the greater thickness of the subcutaneous fat layer, which is more predictive as compared to mouse models when resembling the characteristic of human subcutaneous space [48]. Additionally, larger animals require higher dose of islets to correct diabetes, making it feasible to load them at a high density. *In vivo* studies showed that allogeneic islets encapsulated within the BEAM system successfully reversed diabetes and maintained normoglycemia for over one month and up to 88 days, when experiments were prematurely stopped for logistic reasons (i.e. rats compromising the oxygen transport tubing and electrical wire). These results were achieved at a high cell loading density (60,000 IEQ/mL, equivalent to approximately 4,200 IEQ/cm²). Meanwhile, islets encapsulated in pouches without oxygenation exhibited significant cell death and predominant loss of function. These findings provide proof-of-concept evidence for the feasibility of using oxygenation to maintain islet viability and function at high cell loading densities within poorly vascularized subcutaneous spaces. The loading density of 60,000 IEQ/mL was selected to achieve a reasonable dose (6000 IEQ) of islets to cure diabetes in rats (∼11,000-15,000 IEQ/kg) using a practically small volume of cell suspension (100 μL). While the packing of 60,000 IEQ/mL or 4200 IEQ/cm^2^ is among the highest densities reported in literature (**Supplementary Table 3**), even higher density packing may be possible in clinical application of the BEAM system, especially considering that rat pancreatic islets have a 2-3-fold higher oxygen consumption rate compared to human islets. We estimate that a pouch with an outer diameter of 1 cm and a length of 15 cm would be adequate for encapsulation of approximately 400,000 IEQ at a concentration of 120,000 IEQ/mL (**Supplementary Table 4**). Alternatively, using seven pouches with an outer diameter of 0.75 cm and a length of 7 cm arranged in parallel could achieve the delivery of 420,000 human IEQ at a density of 60,000 IEQ/mL. The final dimensions of the system including the iEOG and connectors would approximate the size of a credit card (**Supplementary Figure 24**). As smaller pancreatic islets (100-150 μm) have been shown to be superior to random-sized pancreatic islets (50-400 μm) in terms of their function [57, 60, 61], the use of smaller sizes may be preferable in preparation of SC-β cell aggregate. We estimate that a system of about half the size of a credit card could be enough to house 500,000 cell aggregates with diameters of 100 - 125 μm. (**Supplementary Figure 25, 26**).

Although the findings from the current design of the BEAM system are promising, several technical challenges must be overcome to realize its potential for clinical translation. Due to the relatively large size of the current electrical controller designed for bench testing, the bioelectronic components could not be fully implanted in rats (a more compact human system has been designed but not implemented). Thus, in this study, we used an extracorporeal setup, wherein, the oxygen generator and electronic components are housed in an external pocket worn by the animal. This setup allows for facile monitoring of iEOG function as well as tracking power and water consumption, which are valuable for the further development of the BEAM system. However, in a clinical context, the entire system—including the cell encapsulation pouch, iEOG, battery, and electrical controller—would need to be fully implantable, with the battery recharged weekly via TET, as is currently used in implantable neurostimulators, left ventricular assist devices and others [62–66]. We acknowledged that using an extracorporeal system, which relies on transcutaneous tubing and an external jacket to secure the electronic components is troublesome, particularly for long-term studies in flexible animals like rats. In fact, experiments were prematurely terminated in 6 out of 9 rats due to damaged oxygen transportation tubing or rats escaping from jackets (**Supplementary Table 2).** Furthermore, although the jackets were initially designed to fit comfortably, the rats could outgrow them within a couple of months due to continuous weight gain as they returned to normoglycemia. The use of larger animals, such as pigs, in conjunction with a fully implantable system, is needed to further assess the long-term efficacy of the BEAM system. Second, we envision that in a fully implantable system, it is necessary to develop a strategy to safely dissipate the hydrogen co-product generated during electrolysis from the body. Although hydrogen has been demonstrated to be safe in other contexts [67–71], such as in scavenging reactive oxygen species, further investigation is required to assess the long-term impact of hydrogen produced by the BEAM system. A study by Zhu et al. demonstrated that hydrogen can be rapidly cleared from rat tissues within 25 minutes, even at saturated levels [72], supporting the feasibility of safely dissipating hydrogen produced by the iEOG. Our simulations indicate that hydrogen generation occurs at a rate of 0.13 scch (5.8 µmol/h) at 280 µA and 4.95 scch (221 µmol/h) at 11 mA. Given the reported clearance rate, these levels are expected to be safely dissipated through an appropriately designed tissue interface facilitating hydrogen diffusion. One potential strategy to enhance hydrogen clearance involves incorporating a porous membrane at the hydrogen outlet, thereby providing sufficient surface area for controlled diffusion of hydrogen in its soluble phase into the surrounding tissue and circulation.

Third, although the accumulation of excessive oxygen around the cell encapsulation pouch did not appear to adversely affect the encapsulated cells, it resulted in a separation of the system from the surrounding tissue in some cases, thereby hindering mass transport. Interestingly, the oxygen production in this study was measured at approximately 2.43 pmol/IEQ/min, a level comparable to or even lower than reported values for rat pancreatic islets in common culture conditions (**Supplementary Table 1**). It is noteworthy that over 90% of the encapsulated islets remained viable and functional, exhibiting responses to glucose changes and maintaining the expression of insulin and glucagon, at the time the excessive gas phase was observed. Given that the oxygen consumption rates of beta cells are variable and highly dependent on environmental factors such as glucose and oxygen levels, it is plausible that the total oxygen produced at a constant rate of 2.43 pmol/IEQ/min would exceed the total oxygen demands of these islets under dynamic conditions. Furthermore, as the implant becomes gradually integrated into the subcutaneous space with newly formed vasculature, the oxygen demand may be less compared to the early post-implantation period. In fact, we observed different levels of FBR and vascularization around the cell encapsulation pouches; thus, the oxygen demands of islets in these implants may vary (**Supplementary Figure 27**). This challenge could potentially be mitigated by incorporating a built-in sensor (eg. total oxygen pressure) within the cell encapsulation pouch, thereby creating a closed-loop system capable of delivering “on-demand” oxygen to the encapsulated cells, as described previously in our patents [73, 74].

In summary, our study underscores the pivotal importance of adequate oxygenation in encapsulation systems to deliver therapeutic cells. Enhanced oxygen levels efficiently sustained the viability and functionality of encapsulated insulin-secreting cells in both *in vitro* and *in vivo* contexts. Notably, continuous oxygen supply allows for the packing of these cells at higher densities without compromising their viability and function. This advancement holds promise for potentially minimizing the required size of encapsulation systems to levels suitable for clinical application, harnessing cell therapy for type 1 diabetes without the need for immunosuppression.

## Experimental Methods

### Preparation of rat INS-1 cell aggregates

INS-1 832/13 cell aggregates were prepared using a commercially available microwell plate (Aggrewell; STEMCELL Technologies). Briefly, 5.9 x 10^6^ INS-1 832/13 cells were added into a well of an AggreWell^TM^ 400 plate, followed by centrifugation at 100 x *g* for 3 minutes. After 48-hour incubation, the cell aggregates were harvested and cultured in Roswell Park Memorial Institute medium (RPMI-1064 GlutaMAX^TM^; Gibco) supplemented with fetal bovine serum (FBS; 10% (v/v); Hyclone), HEPES (10 mM; Gibco), sodium pyruvate (1 mM; Gibco), penicillin (100 IU/mL; Gibco), streptomycin (100 µg/mL; Gibco), and β-mercaptoethanol (50 µM; Gibco).

### Human pancreatic islet

Human islets (Purity 80-95%) were provided by Dr. Chengyang Liu from the Human Islet Core within the University of Pennsylvania, where the islets were isolated according to the guidelines outlined by the Clinical Islet Transplantation (CIT) consortium protocols. These islets were then transported to our laboratory in cold CIT culture media and cultured for 1-3 days at 37°C before encapsulation.

### Rat pancreatic islet isolation

Rat pancreatic islets were isolated from Lewis rats (200-250 grams; Charles River Laboratories) using a previously established enzyme digestion procedure [75–77]. First, 10 mL of Liberase^TM^ TL (0.15%; Research grade, Roche) in ice-cold Hank’s balanced salt solution (HBSS; Corning) was slowly infused into the pancreas through the pancreatic duct. The pancreas was then incised from associated body parts and incubated at 37°C for 31 minutes. Then, 50 mL of ice-cold RPMI-1640 supplemented with 10% FBS was added to terminate the enzymatic digestion. The resulting tissue pellet was washed with RPMI-1640 and filtered through a 450 μm sieve. Islets were purified by a density gradient centrifugation using a Histopaque 1077/RPMI-1640 media gradient. Finally, the islets were manually picked under a light microscope (SZ61, Olympus) to achieve higher purity.

### Preparation and assembly of the BEAM system

#### Manufacture and sterilization of iEOGs

The iEOGs were designed and GMP-manufactured by Giner Inc. Sterilization of the iEOGs was performed using the nitrogen dioxide method (Noxilizer Inc). Before integration with a cell encapsulation pouch, the operational efficacy of the sterilized iEOGs was evaluated using a highly sensitive mass flow meter (Alicat^TM^ Scientific MW-0.5SSCCM-D). Only iEOGs with oxygen production levels within 85% to 115% of their theoretical capability were deemed suitable for use in the experimental procedures.

#### Regulation of oxygen generation

The rate of oxygen generation (Q) is regulated by the modulation of the electrical current (I) applied to the system, as the oxygen flow rate is directly proportional to the current, according to Faraday’s law of electrolysis with adjustments for factors such as temperature, back pressure, and internal losses within the electrolyzer. The equation governing oxygen production is as follows:

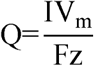

V_m_ is the molar volume of oxygen under standard ambient temperature and pressure (SATP: 25°C and 1 atm). F is Faraday’s constant derived from the product of the elementary charge (e) and Avogadro’s number (N_A_). z indicates the number of electrons required to produce one molecule of oxygen, typically 4 electrons per molecule of oxygen. A custom-built electronic controller was developed by Giner Inc to regulate and maintain a constant current circuit. When the electrolyzer current was set from 0.2 mA to 12 mA, the corresponding voltage required to maintain the set point current was from 1.4 V to 1.8 V, which falls within the expected range for electrolysis.

#### In vitro assessment of iEOG performance

The long-term *in vitro* performance of implantable electrochemical oxygen generators (iEOGs) was evaluated in normal saline. Three iEOGs were configured to operate at a current of 10 mA using a custom electronic controller developed by Giner for 11 month, 2 years, and 2.5 years. Current and corresponding voltage were measured every 30 s throughout the study. On a weekly basis, each unit was connected for an 18-hour period to a sensitive mass flow meter, calibrated at the factory for oxygen measurement (Whisperlite, Model MW-0.5SSCCM-D, Alicat) to measure the oxygen flow rate and demonstrate it corresponded as expected to the current setpoint.

#### In vivo assessment of iEOG performance

To assess the *in vivo* oxygenation capability, sterile iEOGs were implanted into the subcutaneous space of immunodeficient RNU nude rats or immunocompetent SD rats and operated at 11 mA. The flow rates of oxygen generated from these iEOGs were recorded prior to implantation and on the day of explantation. For the experiment on Rowett nude rats, the oxygen generation rates were measured from the iEOGs in tissue samples collected post-mortem. For the experiment on SD rats, a flow meter (Whisperlite, Model MW-0.5SSCCM-D, Alicat) was directly connected to the oxygen outlet of the iEOG exposed through the skin.

#### Preparation of cell encapsulation pouch

The fibrous cell encapsulation pouch was prepared using an electrospinning method. Specifically, medical-grade polyether block amide (Arkema Pebax™) was dissolved in 1,1,1,3,3,3-hexafluoro-2-propanol (HFIP) at a concentration of 12.5% (w/v) and electrospun at 14 kV through a 23-gauge blunt needle, which served as the spinneret. The polymer solution was fed at a flow rate of 1.5 mL/h, with the spinneret-to-mandrel distance set to 11 cm. To fabricate round-ended, edge-free cell encapsulation pouch, a PEG template of the desired shape was deposited onto a conductive mandrel by injection molding to collect electrospun fibers. For the rat scale pouch, a two-part 3D-printed mold was fabricated using a Formlabs 3 printer (Formlabs Inc) with Clear V4 photopolymer resin. The mold was assembled onto one end of a steel rod with a diameter of 1.6 mm, and melted PEG was introduced into central cavity within the mold. After allowing the PEG to solidify at room temperature for 10 minutes, the mold was disassembled, resulting in a 700-μm-thick PEG layer attached to one end of the rod. A 1-cm medical-grade polyurethane ring (Scientific Commodities Inc) was affixed to one end of the PEG template, functioned as a part of the barb connector to seal the pouch. For long pouches, a braided nitinol mesh scaffold (medical grade; Secant Group) with a diameter matching that of the PEG template was first immersed in a 1.5% (w/v) PEBAX solution in HFIP and subsequently positioned over the PEG template. The mandrel was then mounted onto a rotator operating at 450 rpm to facilitate the collection of electrospun fibers for 10 minutes. Afterward, the mandrel was removed and immersed in distilled water for 30 minutes to completely dissolve the PEG template. The resulting pouch was washed 5 times with distilled water and then vacuum-dried overnight. Finally, the pouch was sterilized using an Anprolene AN74 ethylene oxide (EO) gas sterilizer (Andersen Sterilizers) following a standard 12-hour cycle.

The membrane topology was analyzed using a Gemini 500 scanning electron microscope (Zeiss) at the Cornell Center for Materials Research. Fiber sizes were measured with ImageJ from 10 frames taken across 3 independent preparation batches. Tensile tests were conducted using an Instron 5965 test system (Instron) and analyzed with Bluehill 3.0 SOP software.

To prepare the alginate-impregnated cell encapsulation pouch, the one-end-open fibrous pouch described above was first submerged in a 2% SLG100 alginate solution (Novamatrix) for 35 minutes to facilitate the penetration of alginate into the pores of the membrane. Subsequently, a 100 mM barium chloride solution was injected into the pouch through the open end using a 4-inch, 20-gauge blunt needle. The pouch was then held in the alginate solution for 10 seconds to allow the diffusion of barium ions across the fibrous membrane, thereby initiating gelation of the alginate both within and adjacent to the membrane. This process resulted in the formation of a thin alginate hydrogel layer that was securely affixed to the fibrous membrane. The alginate-impregnated pouch was then washed three times with normal saline and stored in a solution containing 95 mM calcium chloride and 5 mM barium chloride until further use.

#### Cell loading and assembly of BEAM system

A 4-cm section of medical-grade PTFE tubing (0.864 mm I.D. x 1.420 mm O.D.; SAI Infusion Technologies), with one end sealed and approximately 1.5 mm perforations along its length, was utilized as a cell loading apparatus. Pancreatic islets or INS-1 cell aggregates were washed three times with normal saline and subsequently suspended in a 2% SLG100 solution. Specifically, for the rat studies, Lewis rat pancreatic islets (6000 IEQ) were suspended in 100 μL of SLG 100 solution. The prepared cell suspension was pre-loaded into another PTFE tubing, which was then connected to the cell loading apparatus centrally positioned within the cell pouch. The other end of the PTFE tubing was connected to a syringe, which was used to gradually infuse the suspension into the cell compartment through the perforations in the loading apparatus. Following this, the cell encapsulation pouch was immersed in a crosslinking solution composed of 95 mM calcium chloride and 5 mM barium chloride for 10 minutes to facilitate the gelation of the cell-embedded alginate layer. The pouch was then washed 3 times with normal saline. The cell loading apparatus was removed, providing space for the subsequent insertion of oxygen transport tubing.

The oxygen transportation tubing is composed of two distinct segments: an oxygen-permeable segment constructed from USP class VI silicone tubing with an I.D. of 0.635 mm and an outer diameter O.D. of 1.194 mm (Specialty Manufacturing Inc), and an oxygen-impermeable segment fabricated from medical-grade polyurethane tubing with an I.D. of 1.016 mm and an O.D. of 1.379 mm (Scientific Commodities Inc). The length of the oxygen-permeable segment was designed to match the length of the cell-laden section of the encapsulation pouch, which was specifically 2 cm in the version used in rats. To mitigate the risk of kinking post-implantation, a miniature coil spring (D.R. Templeman Co.) was inserted into the oxygen-impermeable segment. Furthermore, a polycarbonate barb connector (Bio-Rad) was incorporated onto the oxygen-impermeable tubing. The cell encapsulation pouch was sealed by simply sliding the oxygen transportation tubing fully into the cell capsules, ensuring that the barb connector fits securely within the polyurethane ring of the cell encapsulation pouch. To assess the durability of the connection, a tensile pull test was conducted. The connector and the polyurethane ring were mounted axially in a holding fixture and tested at a rate of 15 mm per min. The force exerted at the moment of separation between the tubing and the barb was recorded. Finally, the sealed encapsulation pouch was connected to the oxygen outlet of an iEOG. All tubing connections were assembled using a custom metal barb connector (Giner Inc), and the junction was reinforced using CA-MG Infinity Bond™ cyanoacrylate adhesive (Infinity Bond).

### Assessment of cell viability

The viability assessment of pancreatic islets and INS-1 cell aggregates was conducted using a dual fluorescent staining technique. Live cells were stained with acridine orange (AO; Sigma Aldrich), while dead cells were stained with ethidium homodimer-1 (EthD-1, ThermoFisher Scientific), following the provided instructions from the supplier. AO/EthD-1 were diluted in RPMI-1640 medium supplemented with 10% FBS, 10 mM HEPES, 1 mM sodium pyruvate, and 100 IU/mL penicillin and 100 μg/mL streptomycin and incubated with islets or cell aggregates for 1 h. The fluorescent images were captured using a digital inverted fluorescent microscope (Invitrogen^TM^ EVOS^TM^ FL, Fisher Scientific).

### Static glucose-stimulated insulin secretion (GSIS)

The functionality of encapsulated pancreatic islets was assessed through a static GSIS assay. Cell encapsulation pouches from three groups—the positive control group (20% oxygen), the negative control group (1% oxygen, without oxygen supplementation), and the experimental group (1% oxygen, with oxygen supplemented from iEOGs)—were subjected to triple rinsing with Krebs-Ringer bicarbonate buffer (KRBB) and preconditioned in a 2.8 mM glucose solution for one hour at 37°C. Subsequently, these pouches were sequentially exposed to a low glucose solution (2.8 mM), followed by high glucose solutions (16.7 mM or 28 mM), and then returned to a low glucose solution (2.8 mM) under 1% O_2_ culture condition, with each exposure period lasting one hour. Insulin concentrations in the supernatants were quantified using an enzyme-linked immunosorbent assay (ELISA) kit (Millipore Sigma). For islets encapsulated in alginate segments retrieved from animal models, the total IEQ count in the segments was determined to normalize the insulin secretion levels.

### Animal model

All animal procedures were approved by the Cornell Institutional Animal Care and Use Committee (IACUC) and supervised by the Cornell Center for Animal Resources and Education (CARE) staffs. Lewis rats (200-400 grams; Inotiv Inc) were used as pancreatic islet donors while SD rats (380-420 grams; Charles River) were used as recipients. The rats were housed within an environmentally regulated room at the Weill Hall Vivarium Animal Facility, following a standard 12-hour dark/light cycle. Daily welfare monitoring was conducted, with specific indicators including marked lethargy, hypothermia, severe dehydration, ataxia, hunched posture, labored breathing, tiptoe or slow ponderous gait, infection at the implant site, and wound dehiscence. Loss of body weight ≥ 20% compared to baseline and BCS (Body Condition Score) < 2 established as a humane endpoint. Several experiments on iEOG performance were conducted at CBSET (Lexington MA) on behalf of Giner Inc under IACUC approvals and similar daily monitoring.

Customized neoprene jackets were supplied by Lomir Biomedical Inc. These jackets were made with exposed flanks and an open abdominal area, a deliberate design to prevent discomfort during extended experiments. The backpack segment was seamlessly integrated into the jacket using Velcro, facilitating easy wearing and removal of the jackets. Rats were acclimated to these jackets for a period of 3 to 14 days before undergoing surgery. Rats wearing jackets were housed individually to prevent potential harm from cage mates. Objects in the cage that could entangle the jackets, such as nest boxes, were removed to avoid incidents Diabetes was chemically induced in SD rats by intraperitoneal administration of pharmaceutical-grade streptozotocin (STZ; Zanosar, Sicor Pharmaceutics). Initially, a freshly prepared STZ solution in 100 mM citrate buffer (pH 4.5) was injected into healthy rats (40 mg/kg), followed by a second dose (20 mg/kg) after 5 days. Blood samples were collected daily from the tail vein starting on day 7 after the first STZ dose to confirm diabetes. Rats with at least seven successive blood glucose measurements exceeding 450 mg/dL were considered diabetic and underwent the implantation procedure.

### Implantation of cell encapsulation pouch

To prepare for cell encapsulation pouch implantation, anesthesia was induced in rats using 4% isoflurane in oxygen and maintained at 2% throughout the surgery. The fur in the right flank was removed using Nair™ cream (Church & Dwight Company Inc), and the surgical field was asepticized with povidone-iodine cotton swabsticks (Dynarex Corp). Then, a 2-cm skin incision was made above the latissimus dorsi muscle. A 5-cm depth subcutaneous pocket was generated parallelly with the ribs using blunt-tip scissors to make space for the implantation of the BEAM system. The cell encapsulation pouch was then introduced into the pocket. A medical-grade polyester tissue cuff (Surgical Mesh^TM^) was integrated where the oxygen transportation tubing exited from the incision to promote healing and prevent infection. The incision was closed and secured to the transportation tubing using a cruciate suture technique. The transportation tubing was finally connected to an iEOG enclosed in a water-refillable chamber placed in the pocket secured on the back of the rat by a customized jacket. The water was replenished on a weekly basis. The overall weight of the system including internal and external components was under 10% of rat body weight.

### Post-implantation assessments

#### Non-fasting blood glucose (NBG) measurement

NBG levels in rats undergoing implantation were assessed at regular intervals using a Contour Next EZ blood glucose monitoring system (Bayer) following BEAM implantation. Graft failure was defined as NBG levels consistently exceeding 250 mg/dL for three consecutive days. *Intraperitoneal glucose tolerance test (IPGTT)*

The metabolic function of transplanted islets in rats was evaluated through an IPGTT conducted on day 30 post-transplantation. After an 18-hour fasting period, rats received an intraperitoneal bolus injection of 50% (w/v) dextrose solution (Phoenix Pharmaceutical Inc) at a dosage of 2.0 g/kg. Blood glucose levels were measured at 0, 15, 30, 45, 60, 90, and 120 minutes following the dextrose injection. Blood samples were collected from the tail vein at 0 and 90 minutes, allowed to equilibrate at room temperature for 30 minutes, and then centrifuged at 2000 ×g for 10 minutes at 4°C to remove clotted material. The serum obtained from the supernatant was stored at -80°C. Serum C-peptide concentrations were quantified using a Rat C-peptide ELISA kit (ALPCO).

#### Assessment of iEOG function on animals

An electronic box (3.5 cm x 3.5 cm x 1.2 cm), containing a battery pack and an electronic controller, was developed by Giner Inc. to regulate the electrolysis and thereby control the oxygen production rate. The system was designed to record data on the function of the electrolyzer and battery levels at five-minute intervals. The box was equipped with a built-in infrared (IR) transmitter for communication with an IR receiver connected to a laptop, facilitating touchless monitoring of iEOG functionality from outside the animal cages. In addition, a light-emitting diode (LED) was incorporated into the system to indicate the status of oxygen production, where green indicates normal production rates and orange signals abnormal production rates. The functionality of iEOGs was assessed twice daily, and data from the electronic box were reviewed every four days to detect any malfunctions related to the electrolysis. The battery was recharged every 4-5 days to maintain continuous operation. Oxygen cessation lasting more than 24 hours was considered the endpoint of an experiment. In such cases, the cell encapsulation pouches were retrieved within 48 hours of the detected oxygen cessation.

#### Ex vivo assessment of encapsulated pancreatic islets post-implantation

The islet-laden alginate hydrogel layer was carefully removed from the retrieved encapsulation pouch and segmented into three portions for subsequent analyses: viability assessment using AO/EthD-1 staining, functionality assessment via GSIS assay, and histological examination. For histological evaluation, the alginate segments were fixed in 10% neutral-buffered formalin solution (Sigma-Aldrich) for 24 hours. To maintain the structural integrity, the hydrogel layer was subsequently embedded in Histogel™ (Thermo Scientific). Samples were then dehydrated, embedded in paraffin (Paraplast, Leica Biosystems), and sectioned at a thickness of 5 µm using a MCT25 manual microtome (Rankin Biomedical). H&E staining was performed utilizing a commercial staining kit (Sigma) in accordance with the manufacturer’s protocol. For immunofluorescence staining, tissue sections were deparaffinized and rehydrated through sequential immersion in xylene, followed by graded ethanol solutions (90%, 75%, 50%, 25%) and distilled water. Antigen retrieval was conducted using a pH 6.0 citrate buffer at boiling temperature, as previously described ^77^. Sections were blocked in PBS containing 1% (w/v) bovine serum albumin (BSA) and 10% (v/v) donkey serum for 2 hours at room temperature to prevent non-specific binding. Subsequently, sections were incubated overnight at 4°C with primary antibodies: rabbit anti-insulin (1:200; ab210560; Abcam) and mouse anti-glucagon (1:200; ab10988; Abcam). Following this, slides were washed five times with PBS and incubated for 1 hour at room temperature with secondary antibodies: Alexa Fluor 594-labeled donkey anti-rabbit IgG (1:400; R37119, Invitrogen) and Alexa Fluor 488-labeled donkey anti-mouse IgG (1:400; A-21202, Invitrogen). After five additional washes with PBS, the slides were mounted using Fluoroshield™ containing 4′,6-diamidino-2-phenylindole (DAPI) (Sigma). Immunofluorescence images were captured using an APEXVIEW APX100 microscope (Olympus).

### Statistical analysis

All statistical analyses were performed using GraphPad Prism 10. Data are presented as individual values, mean ± standard deviation (SD), standard error of the mean (SEM), or box-and-whisker plots, as specified in the figure captions. The Shapiro-Wilk test and F-test were used to assess the normality and variance equality of the data. Statistical significance was determined at a 95% confidence level, with a p-value of less than 0.05 considered significant. For comparisons between two groups, unpaired t-tests were conducted for normally distributed data; otherwise, the Mann-Whitney U test was performed. For comparisons involving more than two groups, one-way ANOVA was utilized. All figures were generated using GraphPad Prism 10, with confidence intervals displayed for relevant data points.

## Supporting information

Supplementary Figure 1. Experimental setup to measure oxygen and hydrogen flow rates generated from an iEOG implanted in subcutaneous space of rats. S

Oxygen generation from the iEOG

Elasticity and shape memory properties of the nitinol scaffold

Elasticity and shape memory properties of the nitinol-reinforced cell encapsulation pouch

## Acknowledgments

We acknowledge the contributions of the staff at the Cornell Center for Animal Resources and Education (CARE) for their expertise in husbandry, clinical care, and suggestions regarding improvements to the animal jackets. We extend our thanks to Giner Inc. for supplying iEOGs and electrical instruments and acknowledge contributions by Giner team members including Thomas Regan, Melissa Schwenk, Griffin Marella, Yamini Mohan, and consultant James Littlefield working with authors Dr. Tempelman and Mr. Stone. We extend thanks to Dr. Chengyang Liu at the Human Islet Core, University of Pennsylvania, for providing human pancreatic islets. We also thank the Histology Lab at the Cornell Animal Health Diagnostic Center for their assistance in processing and preparing histological sections. This work was partially supported by the National Institutes of Health (NIH, 1R01DK105967-01A1; R43/R44DK113536), the Hartwell Foundation, Breakthrough T1D (fka: Juvenile Diabetes Research Foundation) (JDRF, 2-IND-2021-1072-I-X) and Novo Nordisk Company.

## Author contributions

T.T.P., P.L.T., L.T., and M.M. conceived and designed the project. T.T.P., P.L.T., and A.L. prepared the cell encapsulation pouch. T.T.P. and P.L.T. conducted all *in vitro* and *in vivo* cellular studies. P.L.T. performed imaging analysis, ELISA, and tensile test. C.P. measured the oxygen flow rate and captured histological images. S.S. and L.T. designed, supervised and/or conducted iEOG in vitro and in vivo studies. T.T.P. and M.M. wrote the manuscript. M.M. and L.T. supervised the project. J.A.F. discussed the results and provided suggestions on animal surgery. All authors reviewed and edited the manuscript.

## Competing interests

Tung T. Pham, Phuong L. Tran, Linda Tempelman, James A. Flanders, Simon Stone and Minglin Ma are inventors of the technology described in this paper. Tung T. Pham and Simon G. Stone are advisors; Linda Tempelman, James A. Flanders and Minglin Ma are co-founders of Persista Bio Inc, a company formed to commercialize the technology.

