## Supplementary Figure 1. Experimental setup to measure oxygen and hydrogen flow rates generated from an iEOG implanted in subcutaneous space of rats. S for "A Continuously Oxygenated Macroencapsulation System Enables High-Density Packing and Delivery of Insulin-Secreting Cells"

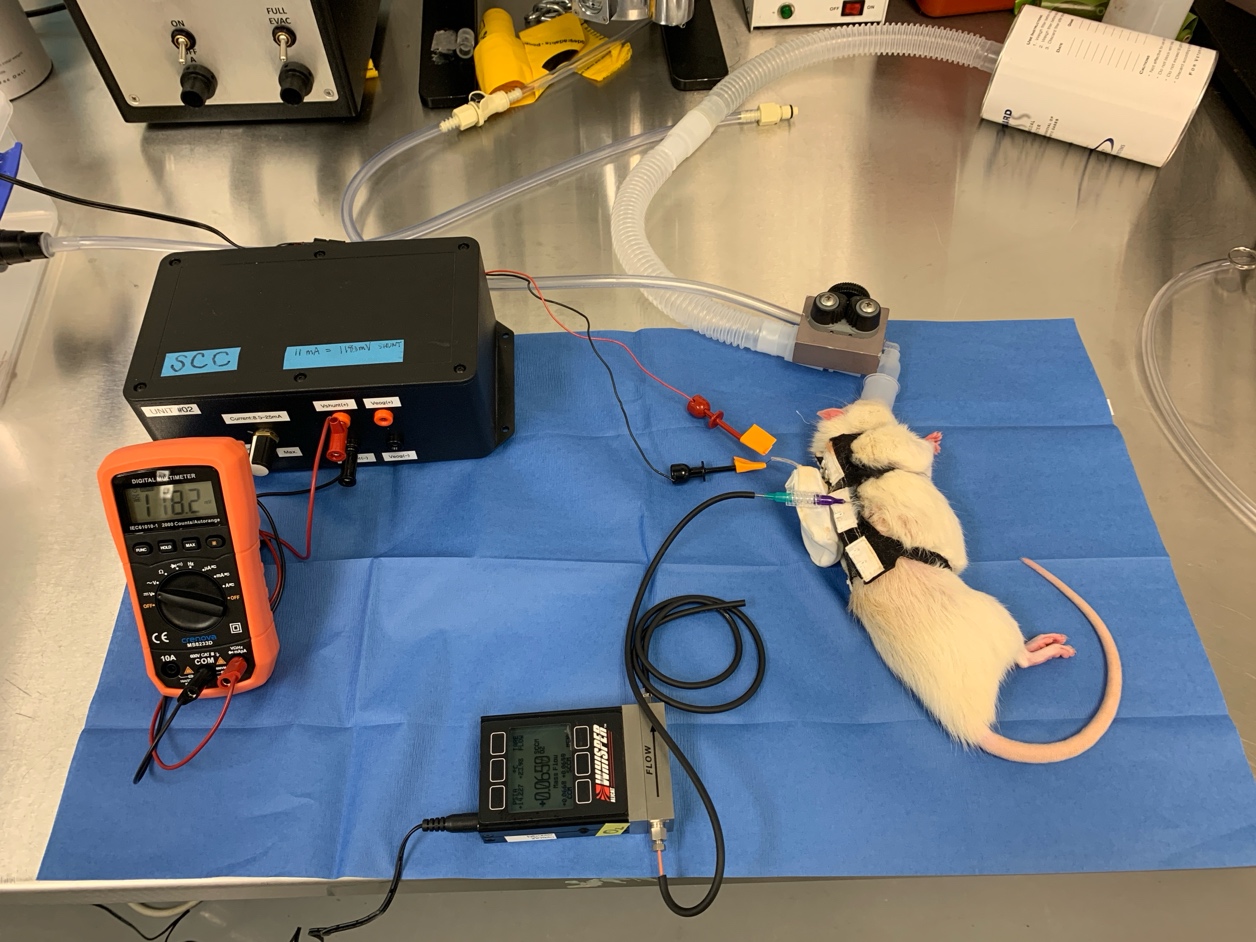

**Supplementary Figure 1**. Experimental setup to measure oxygen and hydrogen flow rates generated from an iEOG implanted in subcutaneous space of rats.

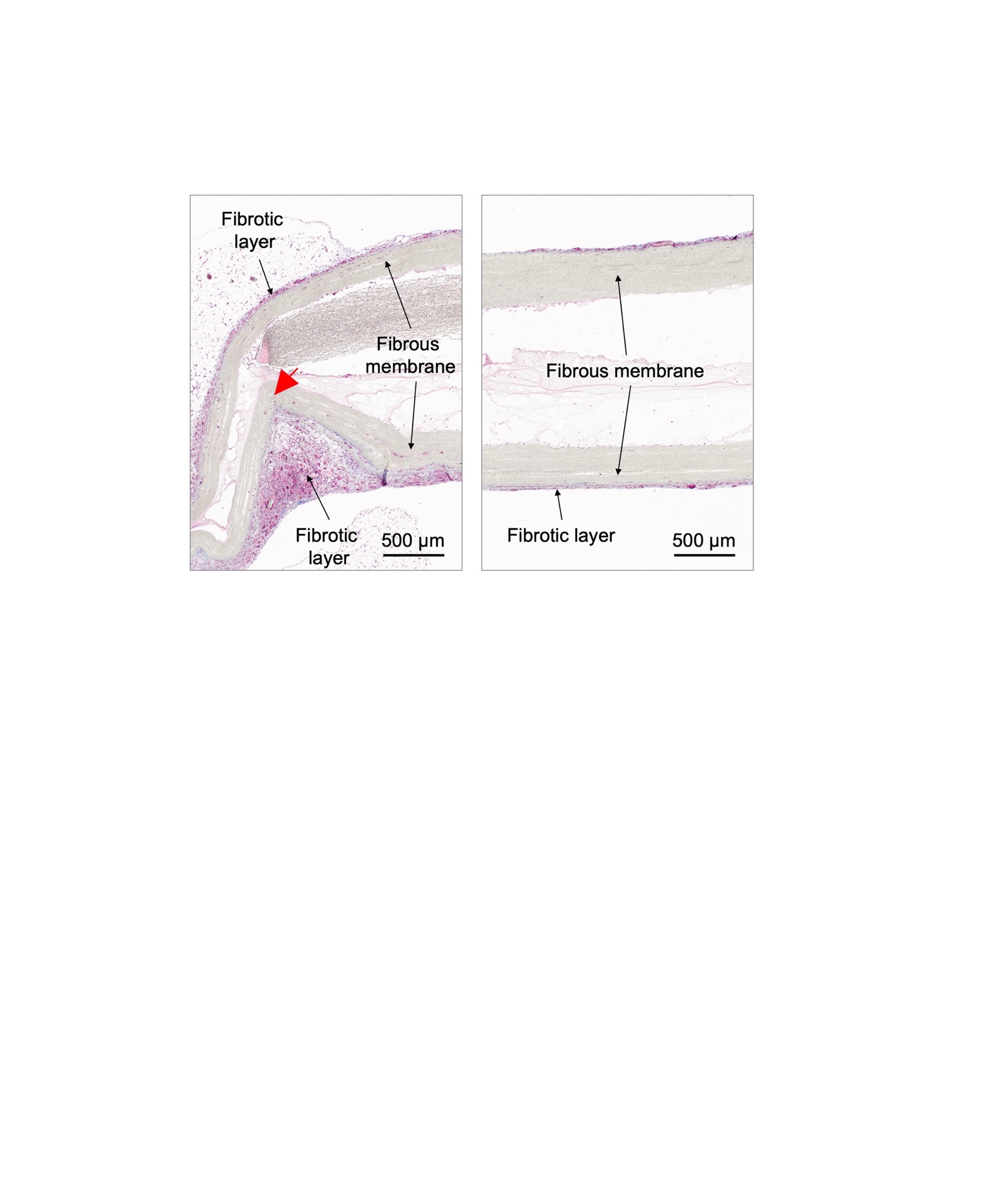

**Supplementary Figure 2**. Masson’s trichrome staining demonstrating different levels of fibrotic response in a kinked region (left panel) and intact membrane (right panel) of a Nylon 6 nanofibrous device implanted into intraperitoneal space of C57BL/6 mice for 1 month. Scale bar: 500 μm.

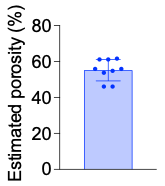

**Supplementary Figure 3**. The estimated porosity of electrospun PEBAX membrane. Data are presented as mean ± SD (n=10). The porosity was estimated using ImageJ software based on 10 SEM images taken from 3 independent batches.

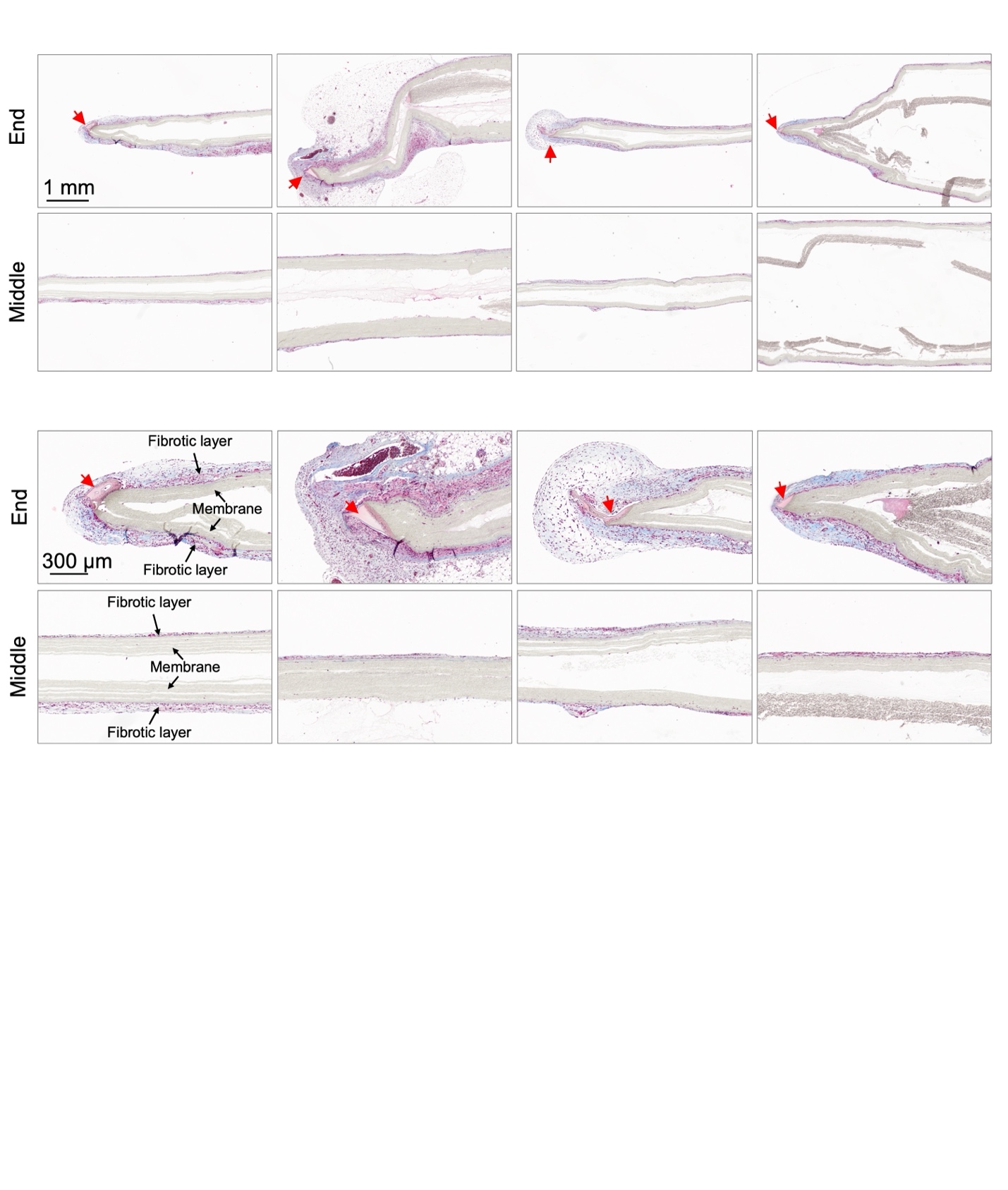

**Supplementary Figure 4**. Masson’s trichrome staining of heat-sealed nanofibrous nylon 6 devices implanted in the intraperitoneal space of C57BL/6 mice for 1 month, demonstrating different levels of fibrotic reactions at the heat-sealed end and the middle of the devices (n=4). Scale bar: 1 mm (upper) and 300 μm (lower). Red arrowheads indicate heat-sealed regions.

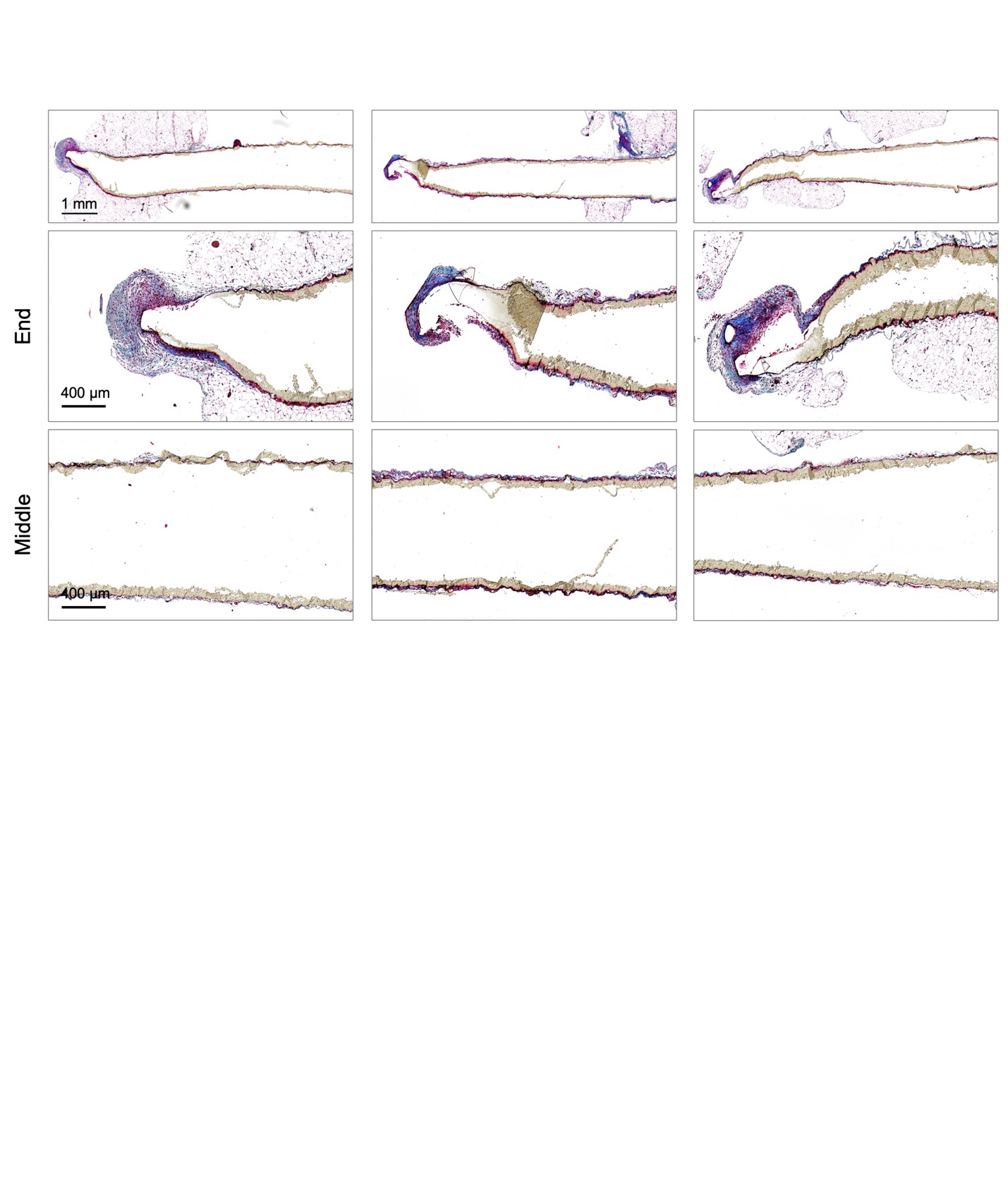

**Supplementary Figure 5**. Masson’s trichrome staining of heat-sealed nanofibrous polyurethane-polyether devices implanted in the intraperitoneal space of C57BL/6 mice for 1 month, demonstrating different levels of fibrotic reactions at the heat-sealed end and the middle of the devices (n=3). Scale bar: 1 mm (upper) and 400 μm (lower).

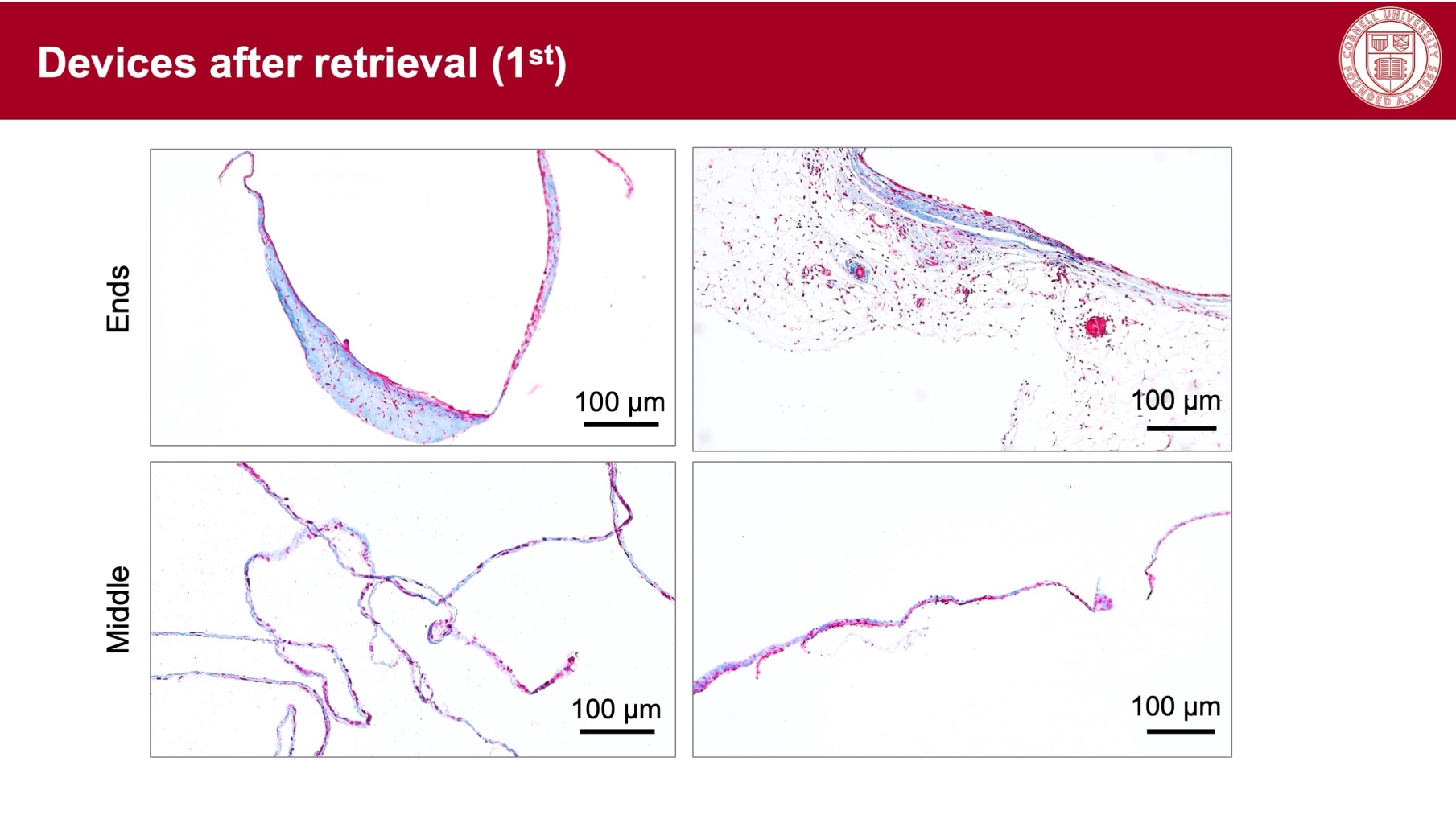

**Supplementary Figure 6**. Masson’s trichrome staining of heat-sealed nanofibrous PEBAX devices implanted in the intraperitoneal space of C57BL/6 mice for 1 month, illustrating different levels of fibrotic reactions at the heat-sealed end and the middle of the devices. Scale bar: 100 μm. The membrane was dissolved during the histology process.

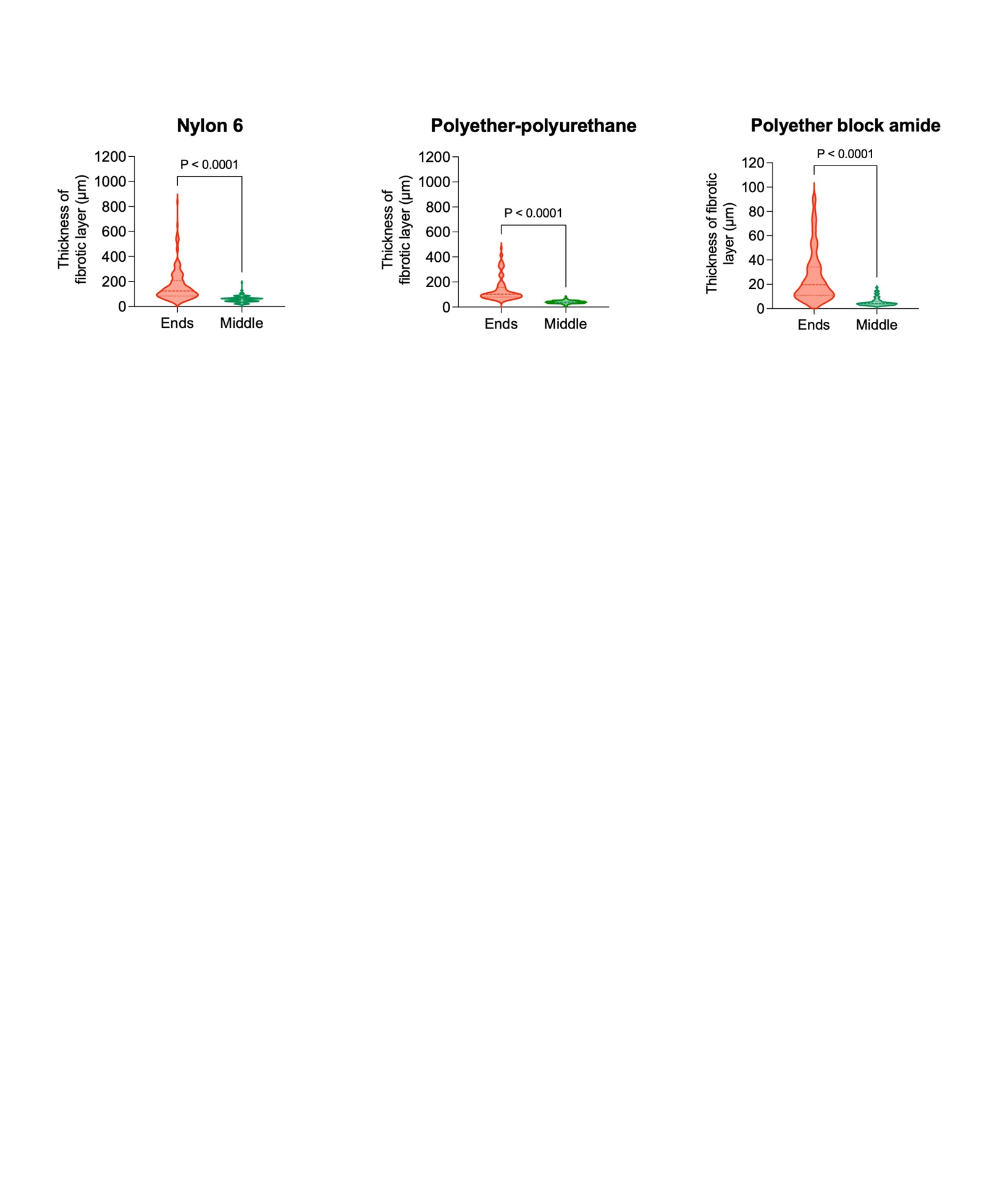

**Supplementary Figure 7**. The thickness of the fibrotic layers at the end and in center of heat-sealed nanofibrous devices made of Nylon 6 (from 4 devices), polyurethane-polyether (from 3 devices), PEBAX (from 4 devices). The data are presented as violin plots. Statistical analyses were performed using unpaired two-tailed Student’s t-test.

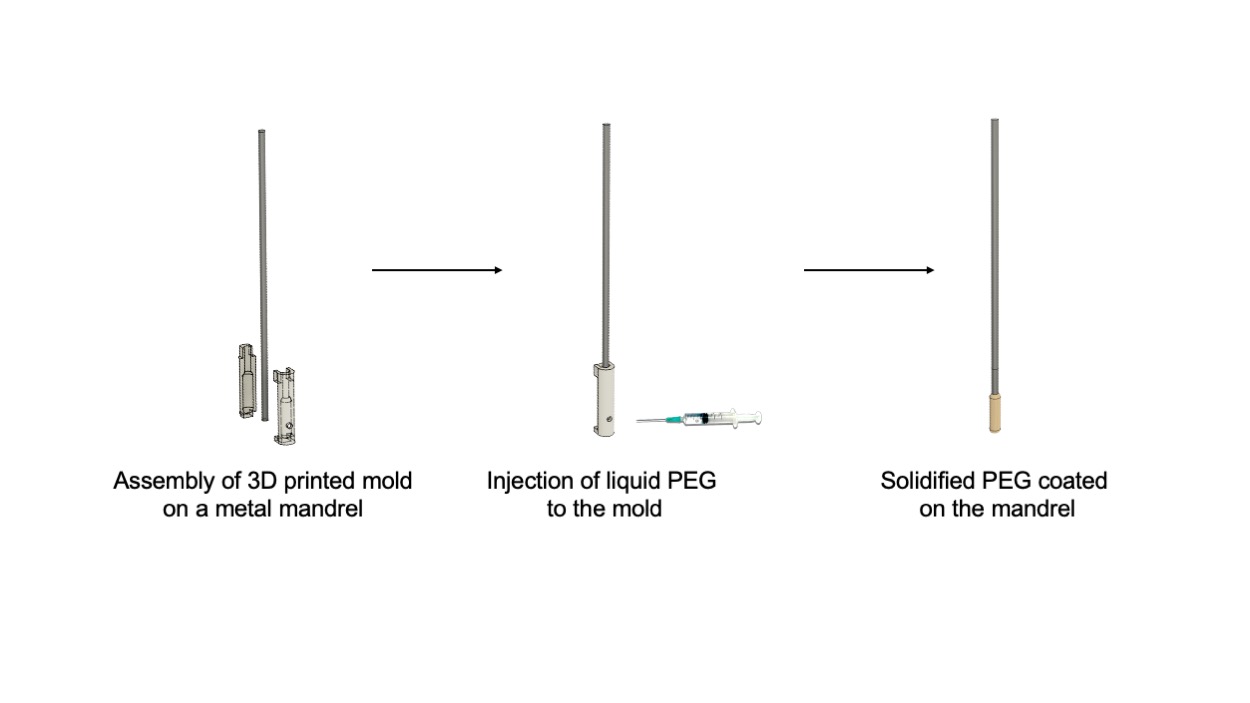

**Supplementary Figure 8**. Schematic illustration for preparation of polyethylene glycol template to collect electrospun fibers.

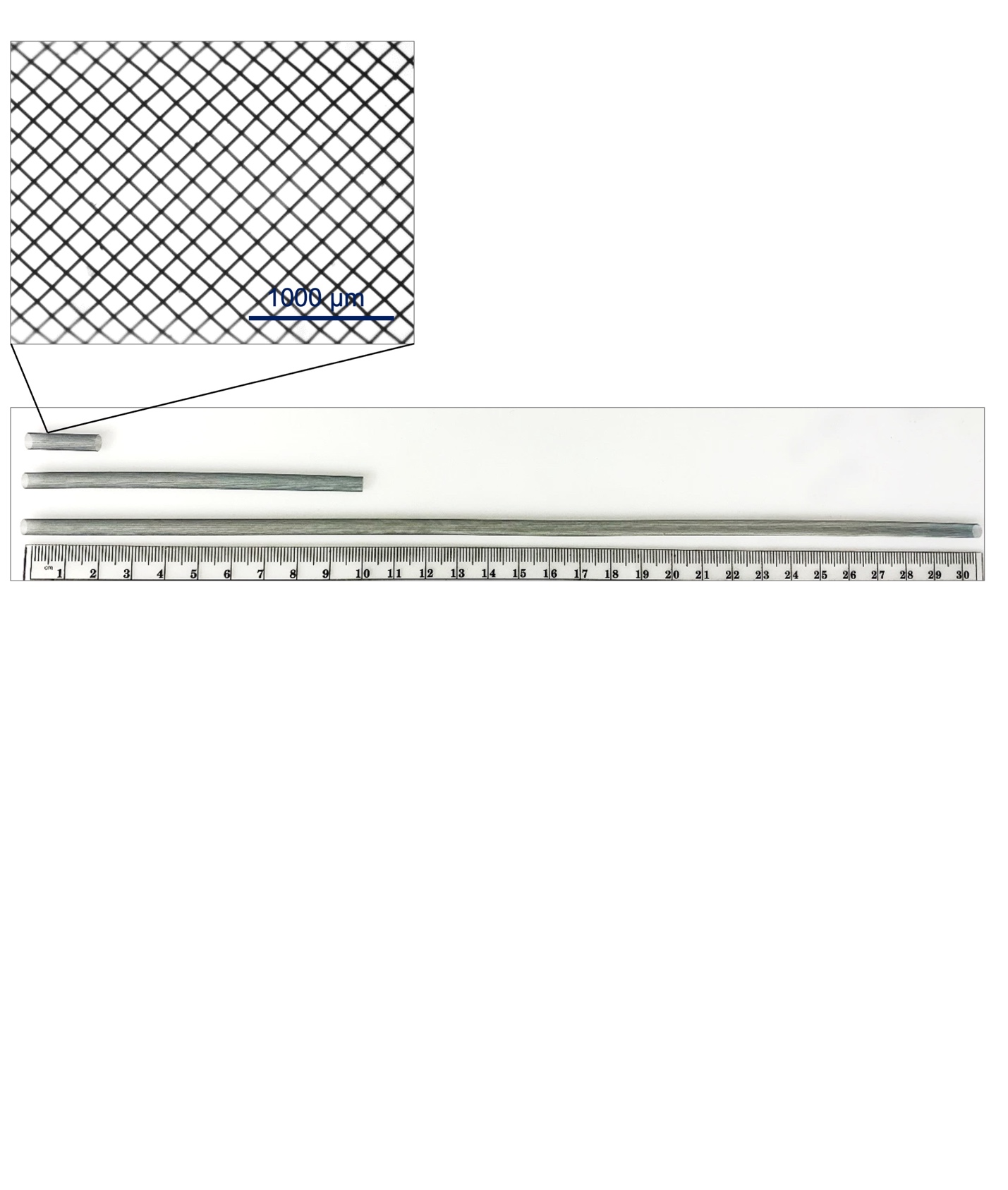

**Supplementary Figure 9**. **Structure of the nitinol braided mesh scaffolds**. The scaffold was constructed from 25.4-μm nitinol braided wires, with pore sizes ranging from 100 to 150 μm. These elastic, shape-memory scaffolds effectively maintained the structural integrity of the immuno-protective membrane while minimally impacting mass transport due to their large pore sizes and thin wire thickness.

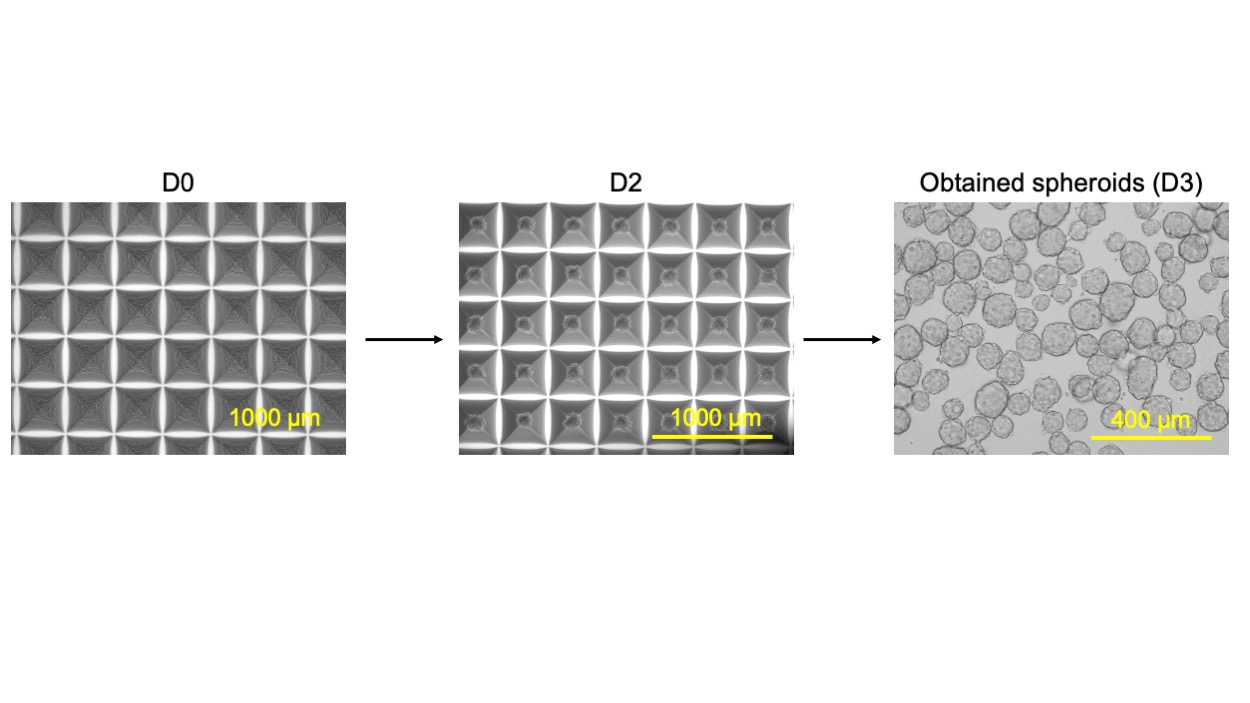

**Supplementary Figure 10**. Preparation of INS-1 spheroids using Aggrewell^TM^.

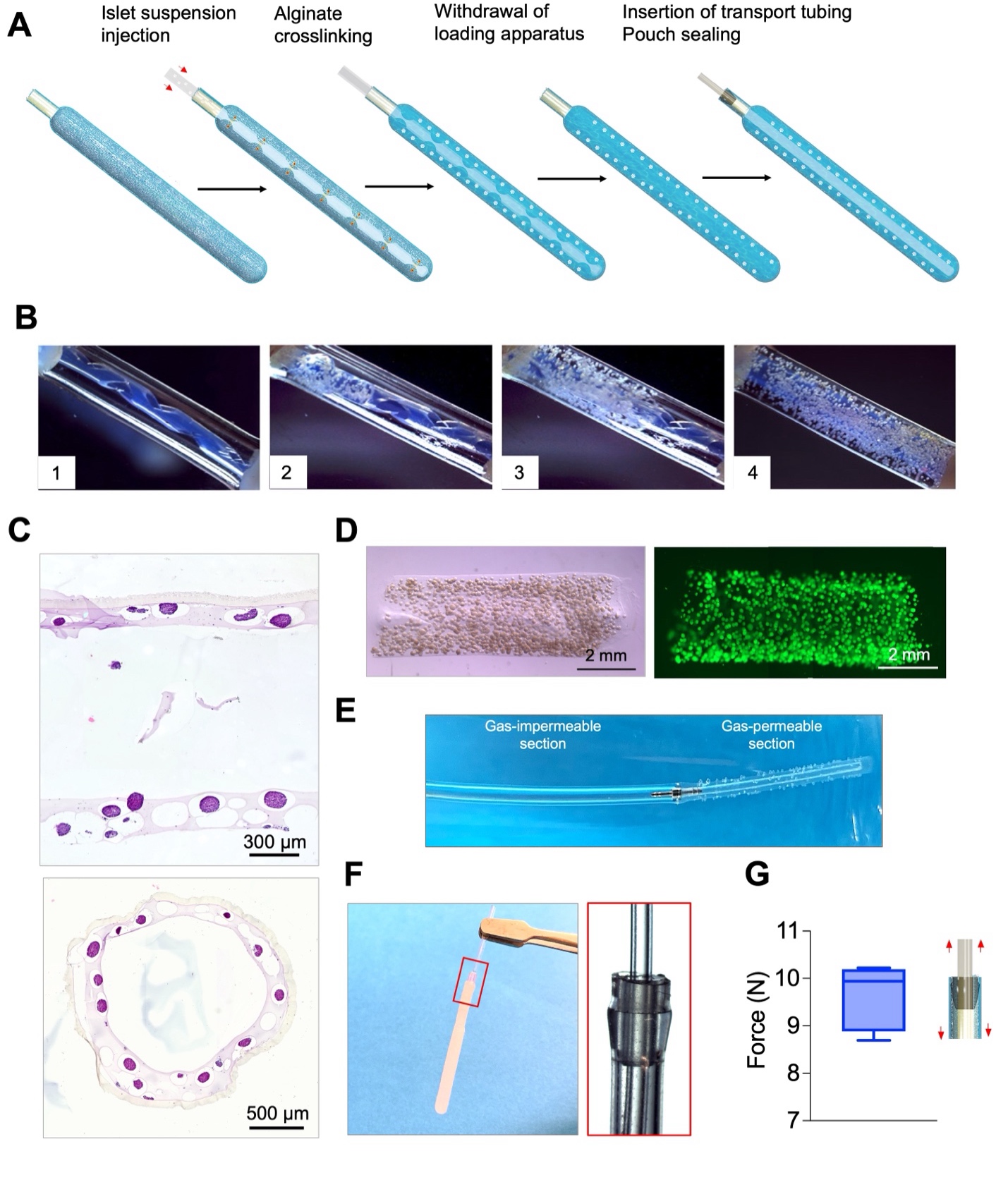

**Supplementary Figure 11**. Macroscopic images showing the distribution of 150-μm beads, used as islet mimics, during the injection of suspension into the encapsulation pouch. Gradual dispersion of beads through the pores of the loading tool results in the occupation of interstitial spaces between the loading tool and the encapsulation pouch wall (a transparent tube used for visualization purpose).

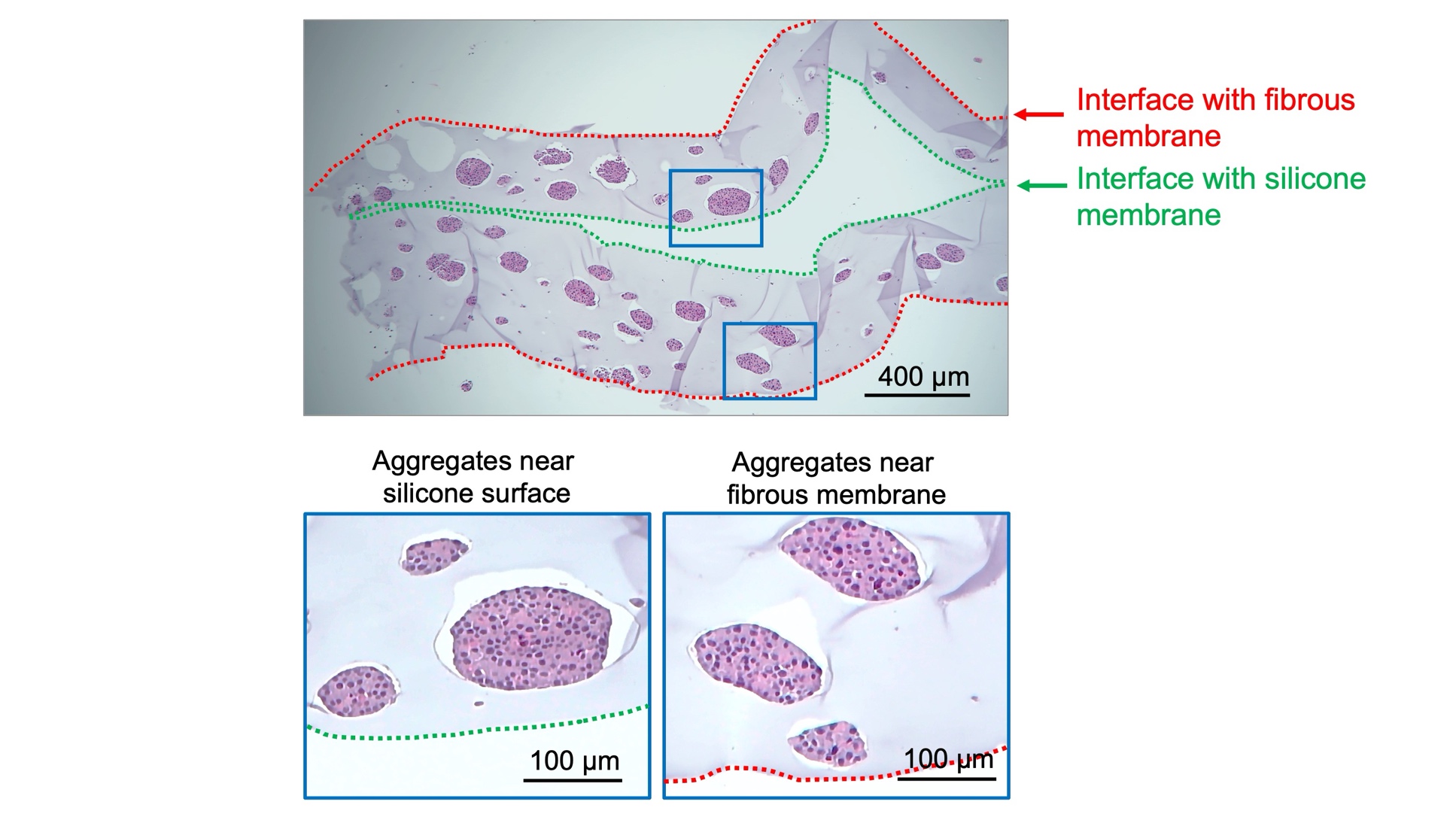

**Supplementary Figure 12**. **A longitudinal H&E-stained histological section of a BEAM system encapsulating INS-1 cell aggregates at a density of 60,000 IEQ/mL**. The system was subjected to 1% O_2_ culture condition for 24 h while the iEOG was operated at 0.27 mA Scale bars: 400 µm (upper) and 100 µm (lower). The sample was sectioned through the silicone core. No significant difference in the viability were observed between cell aggregates located proximal and distal to the oxygen source.

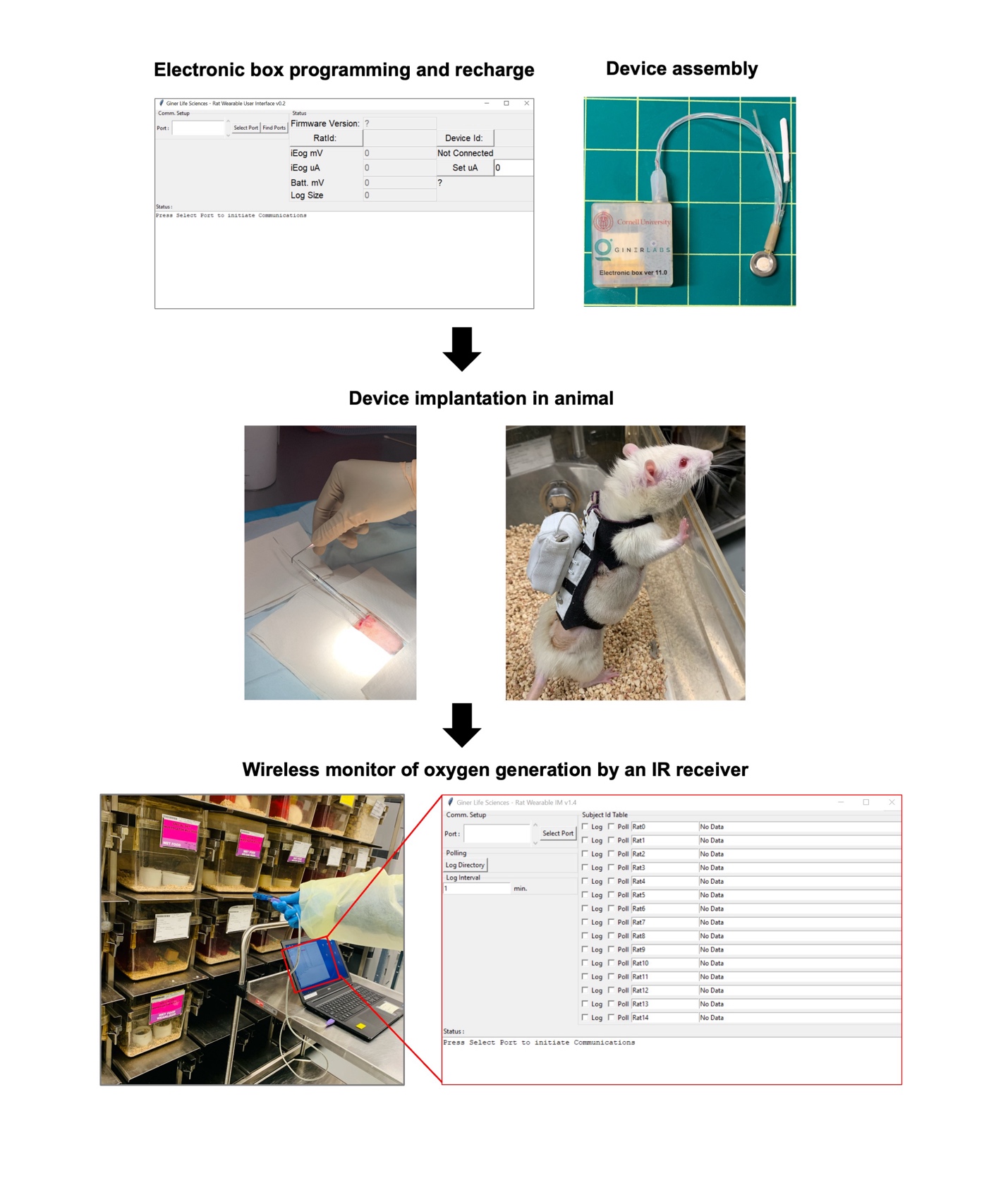

**Supplementary Figure 13**. Workflow in implantation of BEAM device in rats. The electronic controller was programmed to regulate the electrical current, generating an adequate level of oxygen for the islet dose. The electronic controller was then connected to the iEOG on the back of the rats. iEOG performance was wirelessly monitored from outside of the animal cages by an IR receiver linked to a laptop.

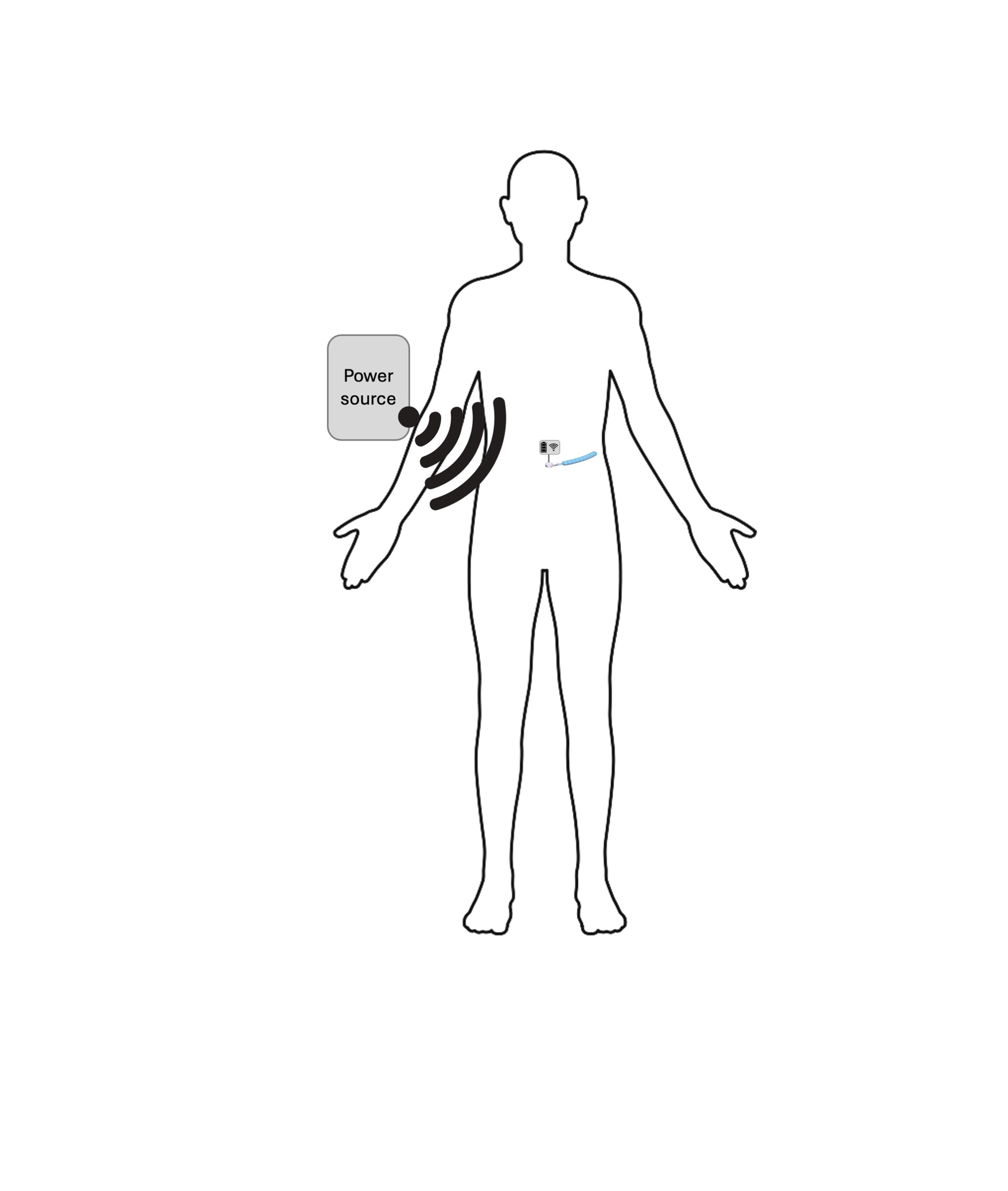

**Supplementary Figure 14**. A proposed setup for the use of transcutaneous energy transfer system for battery recharge of the BEAM device.

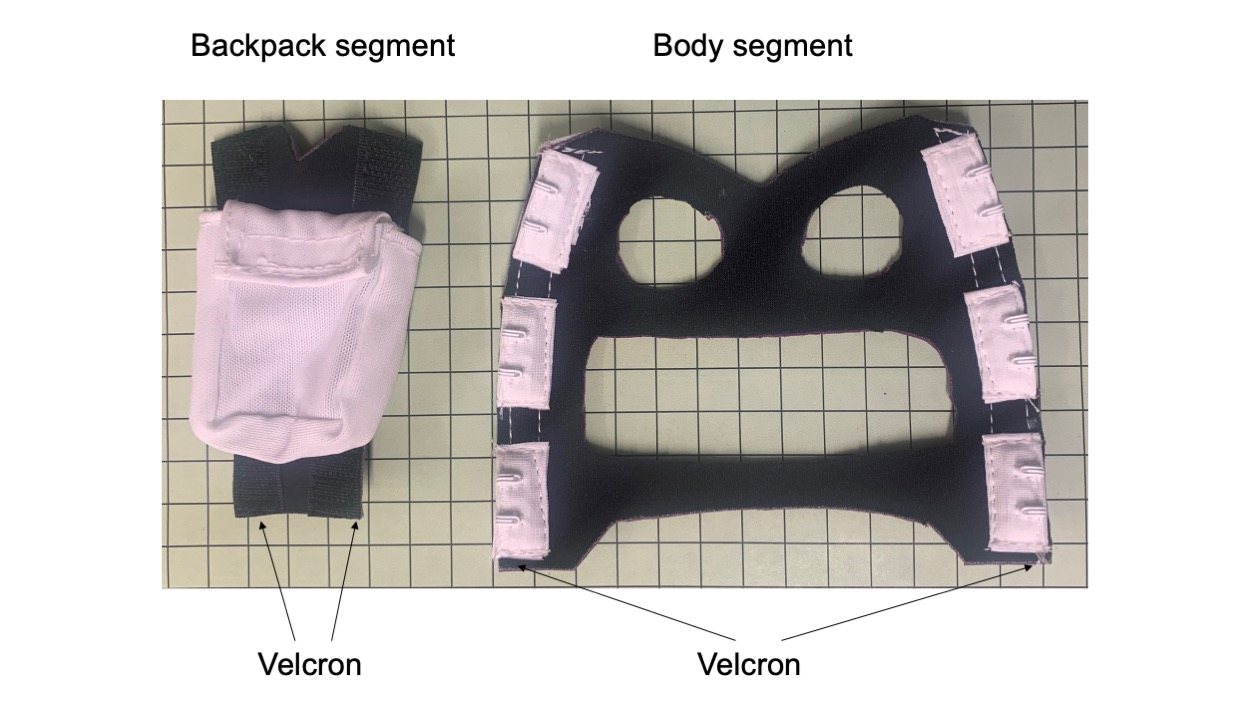

**Supplementary Figure 15**. A digital image of a rat jacket designed with open flank and abdomen to avoid discomfort.

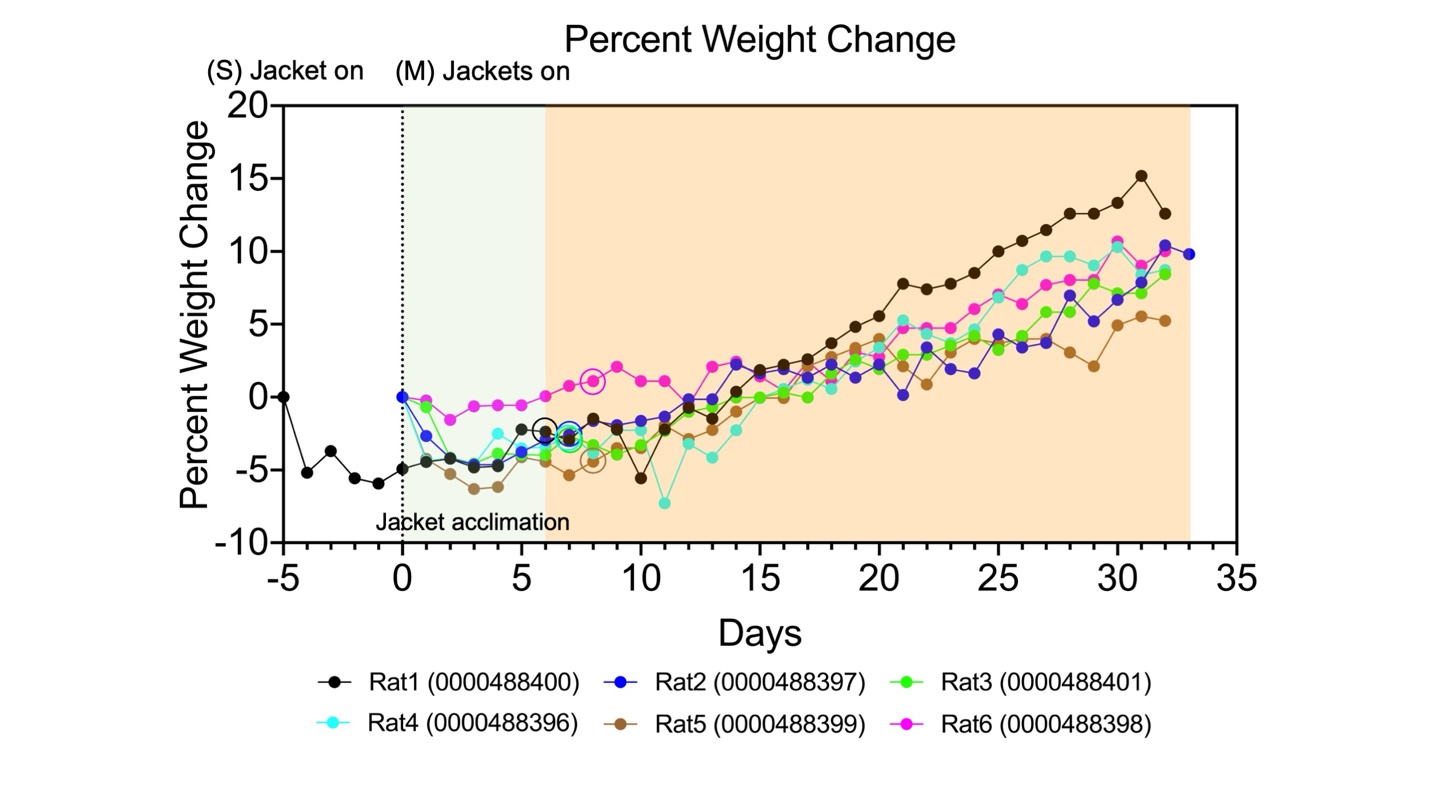

**Supplementary Figure 16**. Percentage body weight change of rats after wearing jackets. The circles indicate the time points that the electronic controllers were placed into the pocket on the back. (n=6).

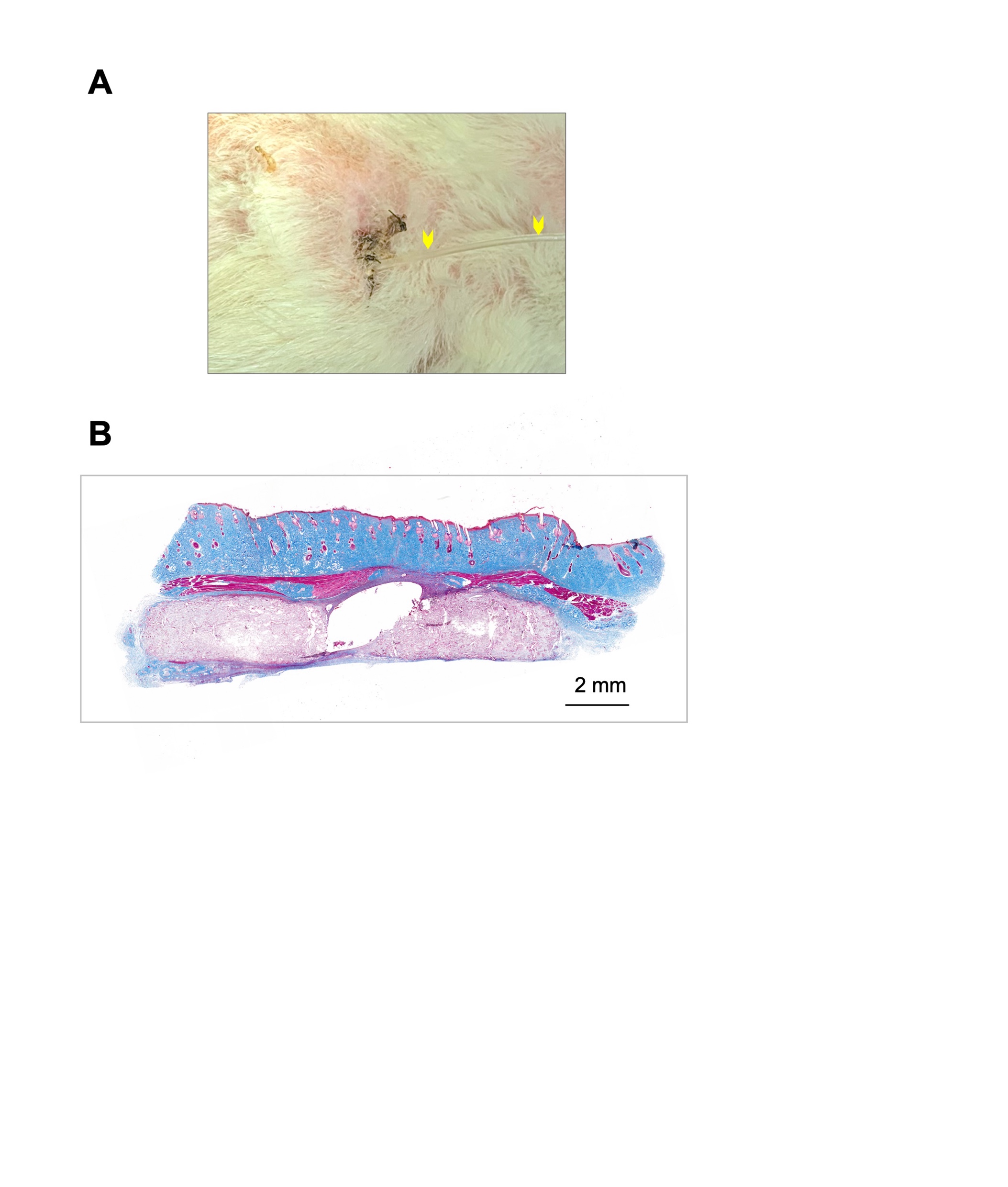

**Supplementary Figure 17**. Assessment of transcutaneous exit after implantation of BEAM device. (**A**) A digital image showing the tissue healing at the transcutaneous exit. Arrows depict the oxygen transportation tubing. (**B**) Masson’s trichrome staining of the transcutaneous exit, showing tissue integration into the polyester cuffs. No adverse reactions or inflammation were observed surrounding the oxygen transportation tubing. Scale bar: 2 mm.

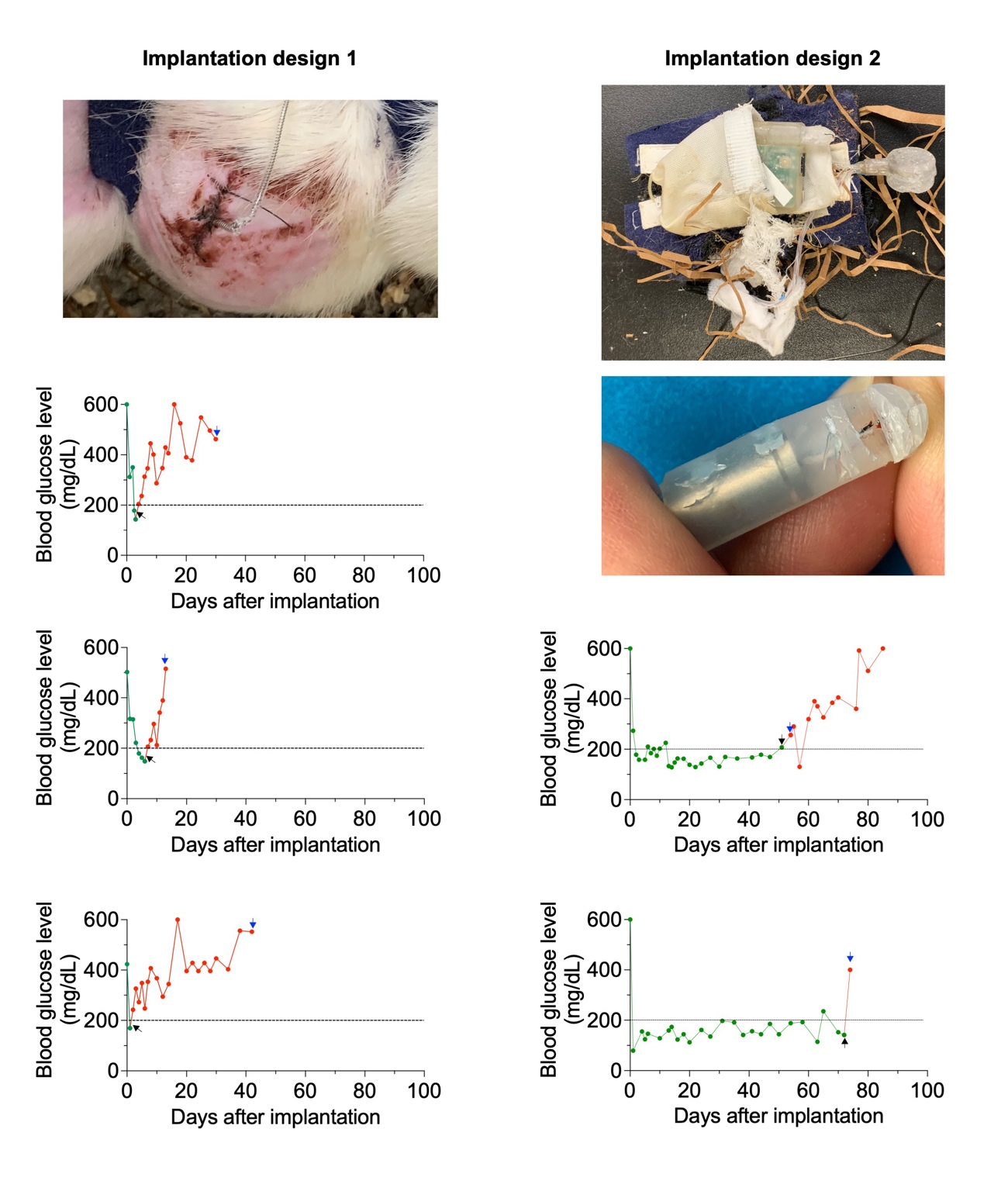

**Supplementary Figure 18**. Premature oxygen cessation caused by rats. For implantation design 1, oxygen transportation tubing was damaged by the rats (3/3), leading to oxygen leakage. For implantation design 2, rats (2/6) escaped from jackets and damaged the electronic components, leading to oxygen cessation for over 24 hours. The black arrows indicate the time points of oxygen cessation while blue arrows mark the time points of device retrieval.

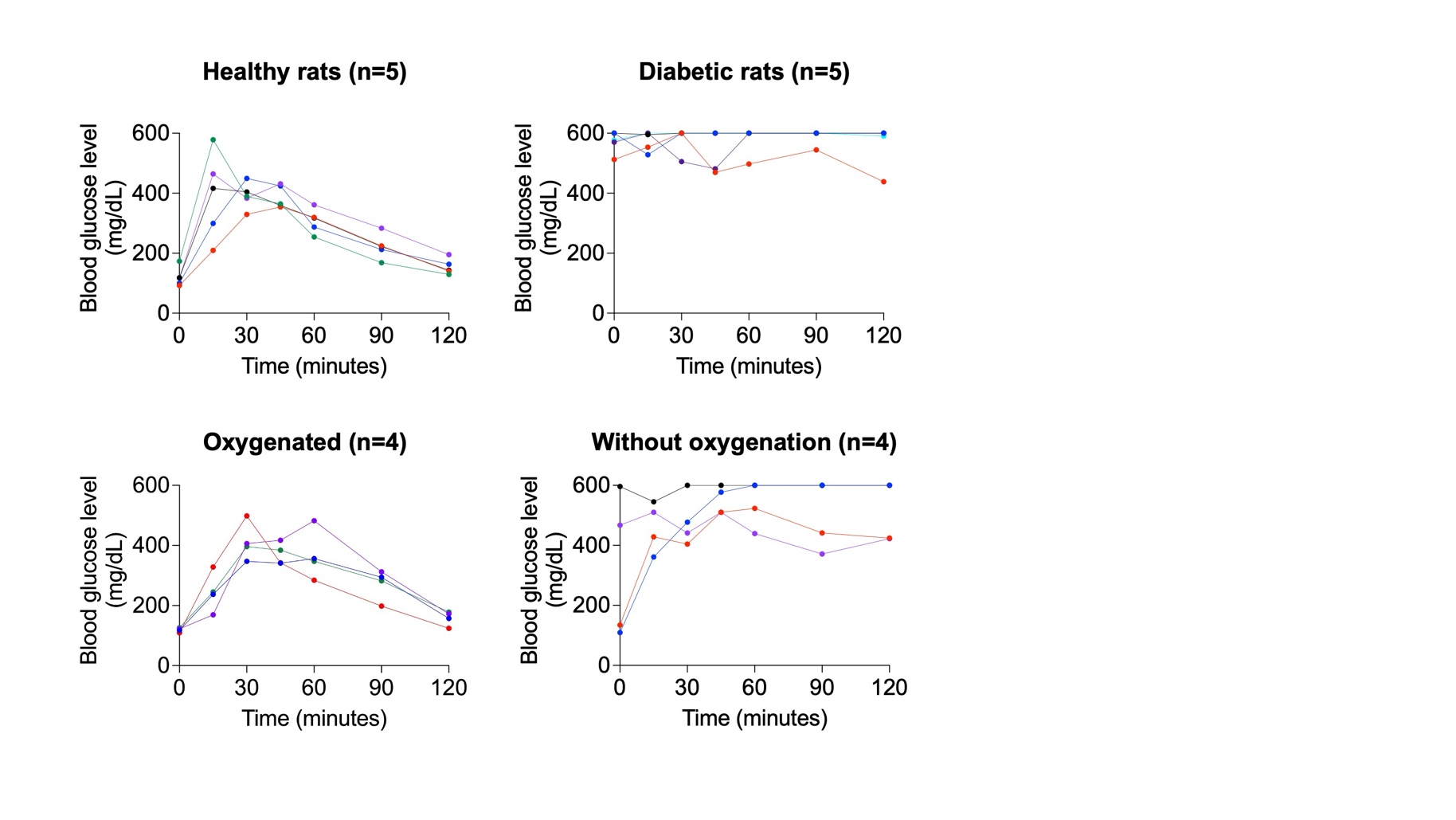

**Supplementary Figure 19.** Intraperitoneal glucose tolerance test conducted on healthy rats (n=5), diabetic rats (n=5), rats receiving a BEAM system (n=4; measured on day 35), and rats receiving a cell encapsulation pouch without oxygen supplementary (n=4; measured on day 35). Data are shown for individual subjects.

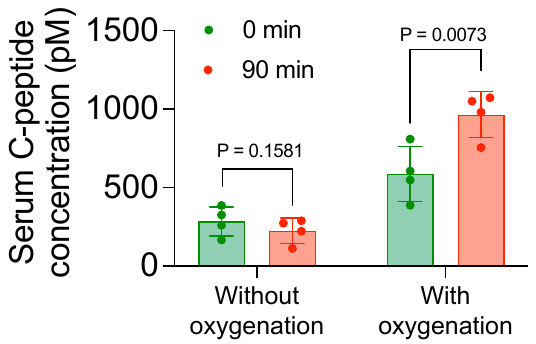

**Supplementary Figure 20.** Serum C-peptide analysis of rats receiving devices without oxygenation (n=4) and with oxygenation (n=4) on day 35 post-transplantation before and 90 min after glucose administration. Non-significant differences were found between C-peptide levels of rats receive non-oxygenated devices before and 90 min after glucose administration. Higher levels and significant increases in serum C-peptides were observed in rats received oxygenated devices after glucose administration. Statistical analyses were performed using paired two-tailed Student’s t test.

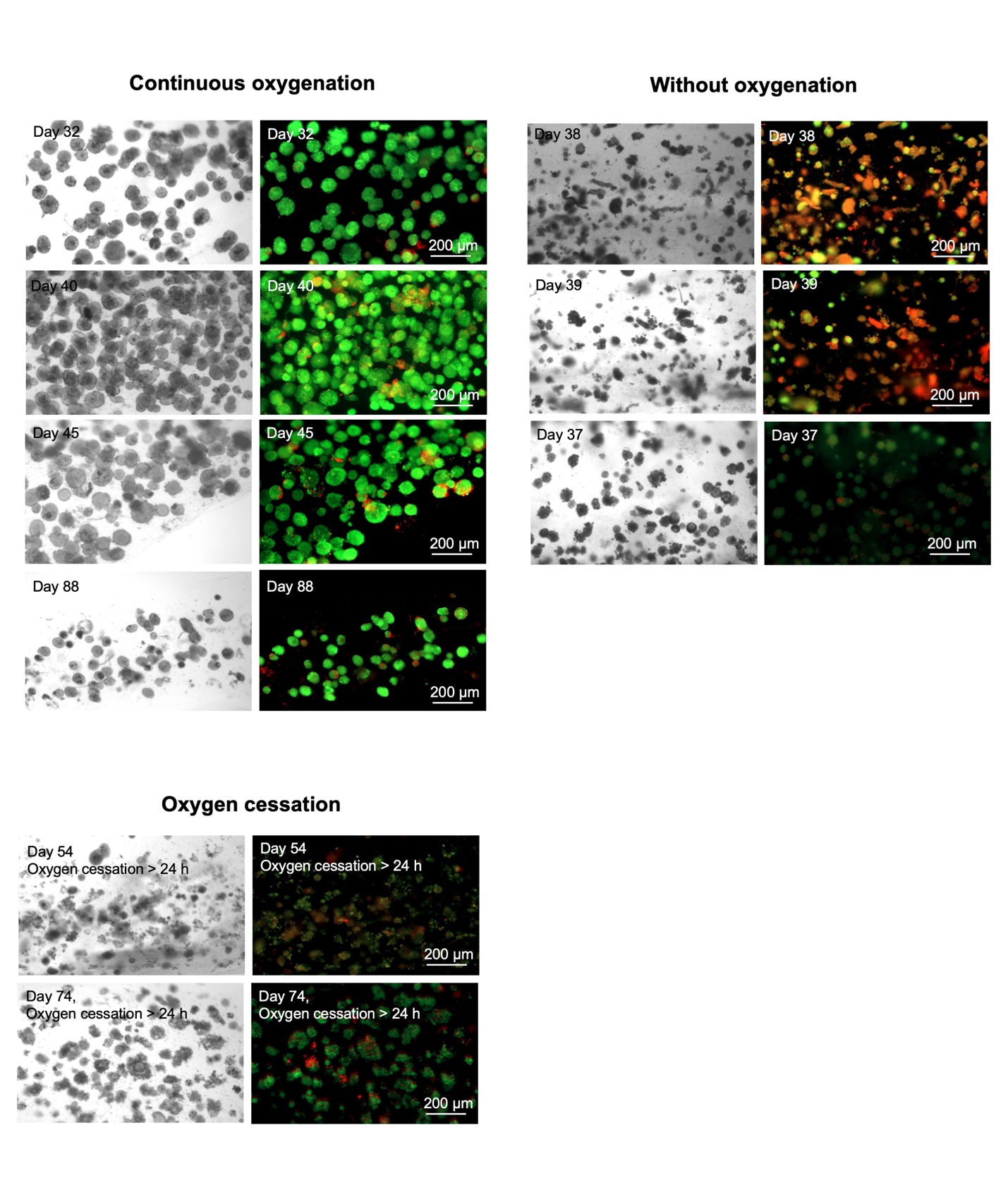

**Supplementary Figure 21**. Assessment of islet viability in devices without supplemental oxygenation, with continuous oxygenation, and with oxygen cessation > 24 hours by dual fluorescence staining. Green: Live cells; Red: Dead cells. Scale bar: 200 μm.
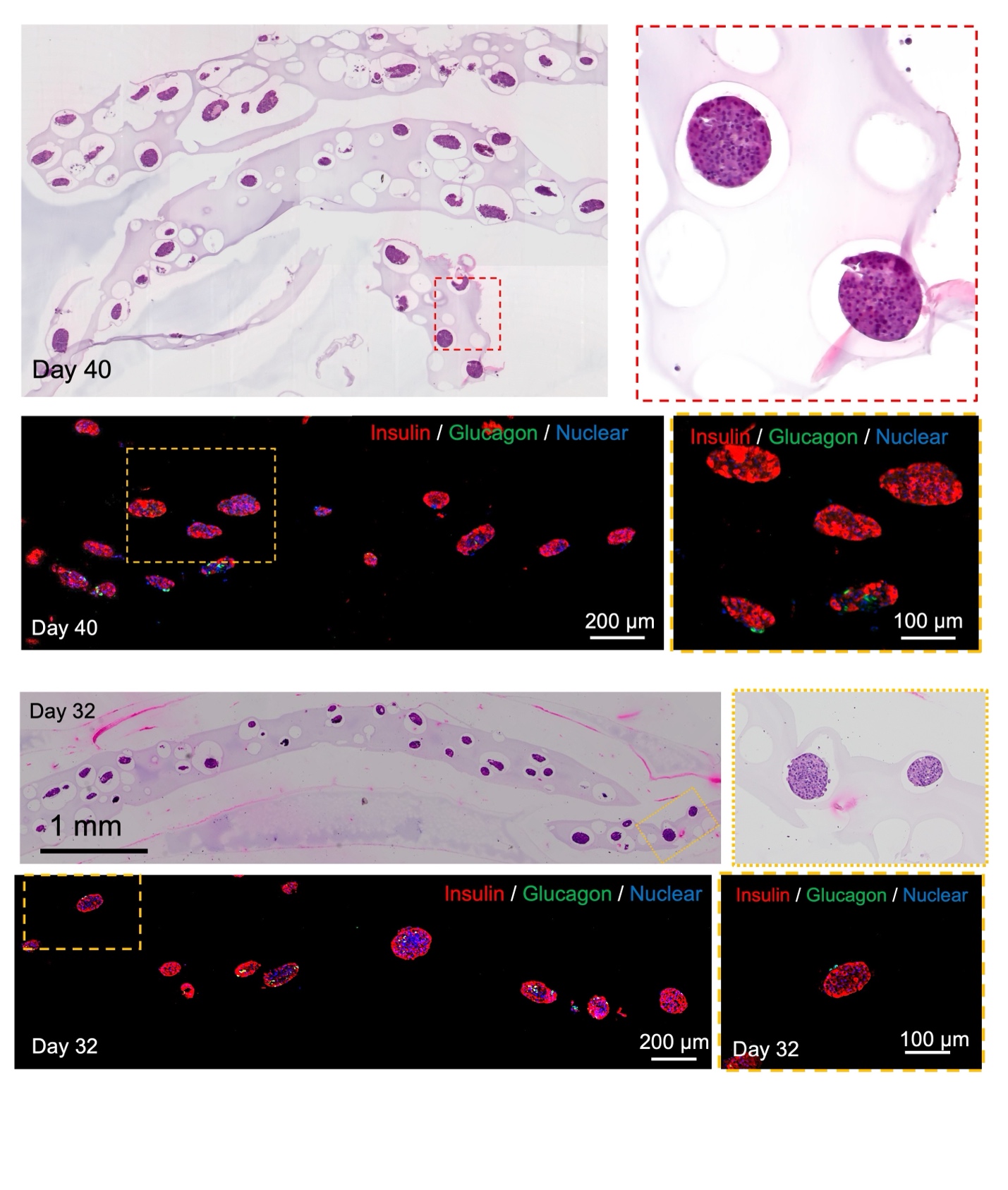

**Supplementary Figure 22**. Representative H&E staining and immunofluorescence staining of encapsulated islets in implants with oxygenation retrieved on day 32 and day 40 due to the formation of gas phase. Red: Insulin; Green: Glucagon; Blue: Nuclear staining.

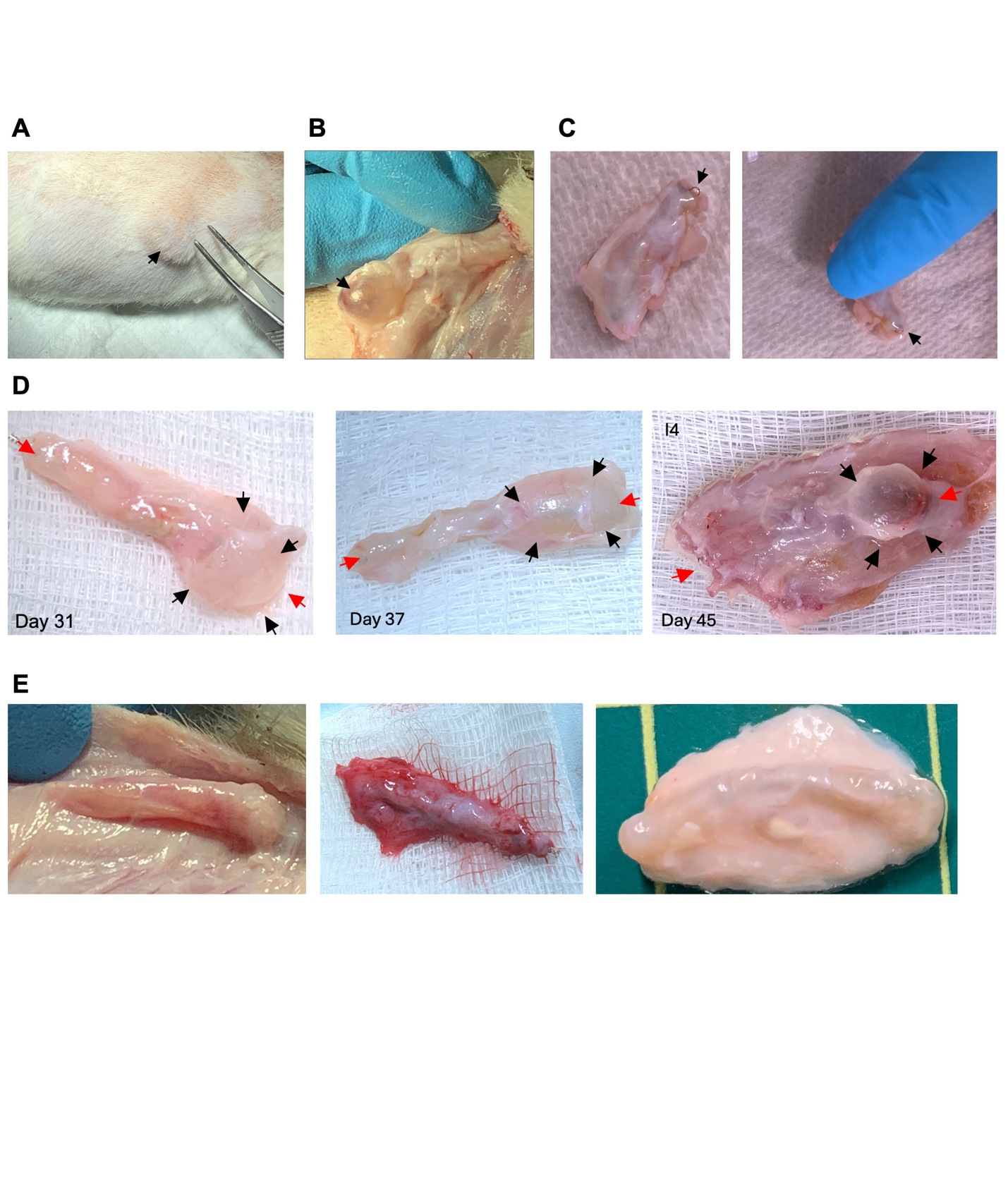

**Supplementary Figure 23. Formation of excessive gas surrounding the cell encapsulation pouch. (A)** Digital image showing the location of the gas bubble at the implantation site. (**B-D**) Gas accumulation surrounding devices retrieved from rats with elevated blood glucose levels. The bubbles were released from the tissue upon device retrieval. Black arrows denote gas bubbles, while red arrows indicate two ends of the cell encapsulation pouch. (**E**) Absence of gas phase in devices with oxygen cessation.

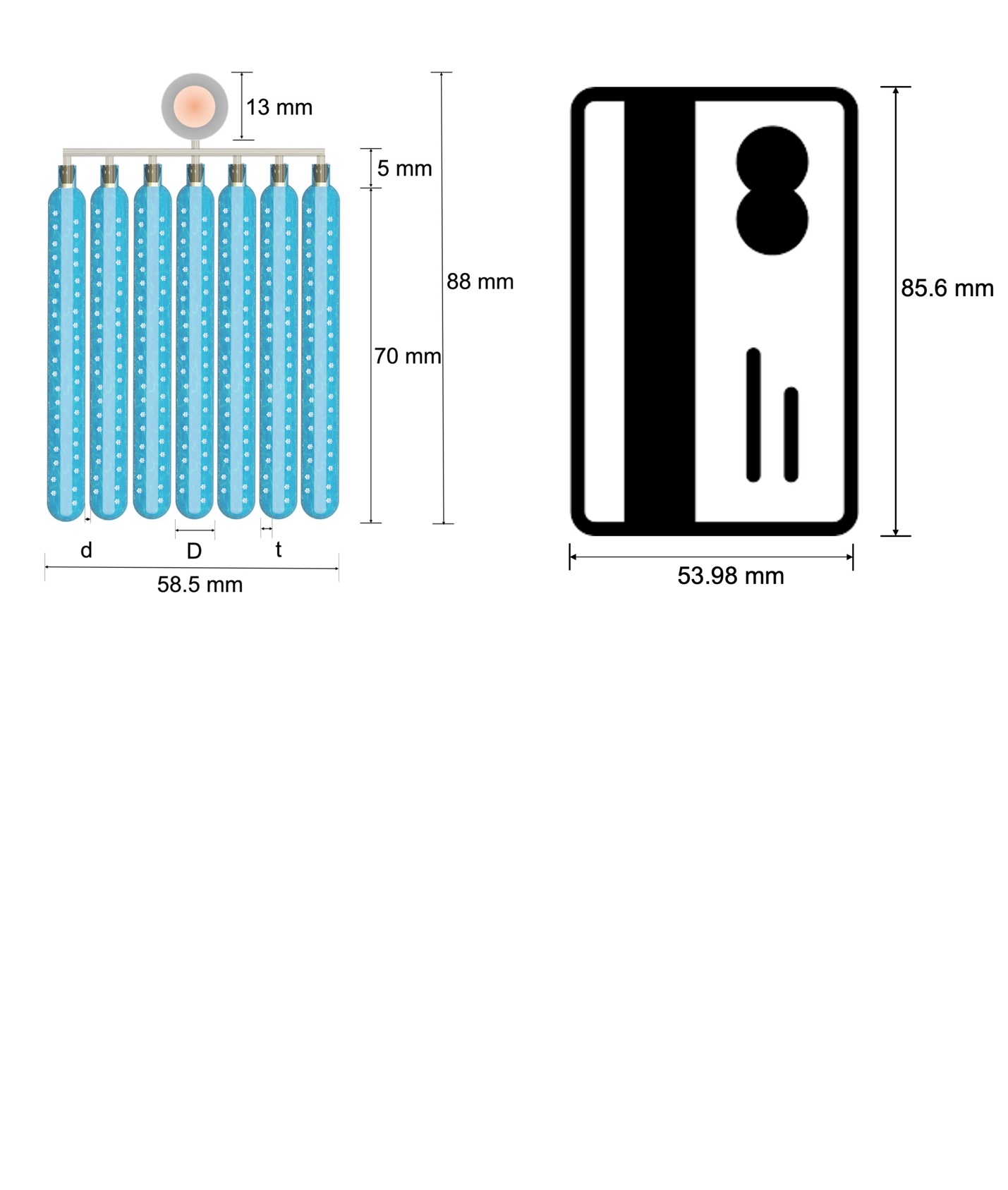

**Supplementary Figure 24**. A proposed method for scaling up the BEAM device aims to accommodate 420,000 human IEQ at a loading density of 60,000 IEQ/mL. Seven cell pouches, each measuring 70 mm in length and 7.5 mm in diameter (D) with a capacity of 1.11 mL per pouch, can be arranged in parallel to deliver the required 420,000 human IEQ. The distance (d) between cell pouches is 1 mm. The thickness of cell-laden alginate hydrogel layer (t) is 0.75 mm. The final dimensions of the device, including the iEOG and connectors, are approximately the size of a credit card. A smaller design could be achieved by increasing the loading density, as human islets consume 2-3 times less oxygen compared to rat islets.

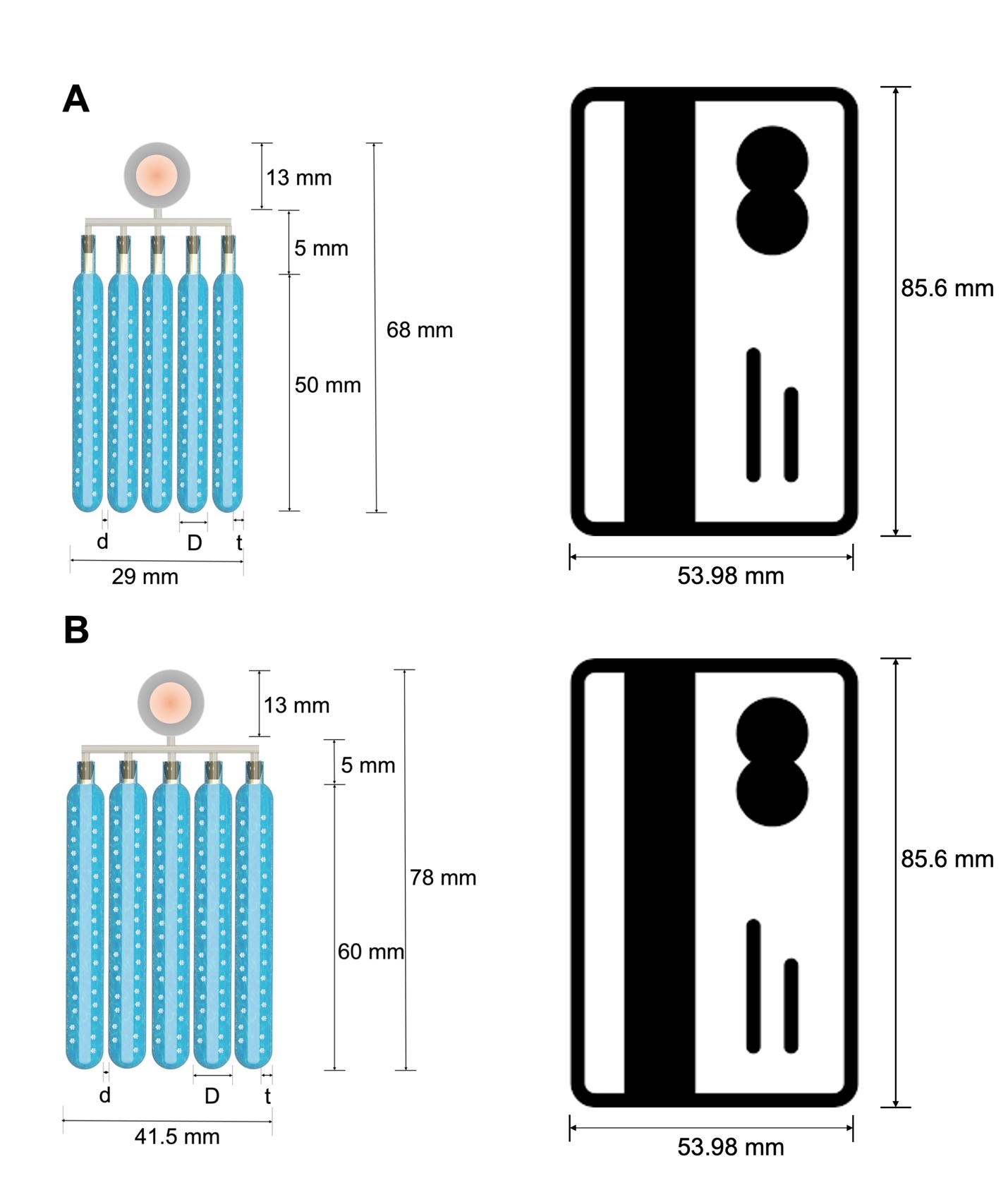

**Supplementary Figure 25.** A proposed method for scaling up the BEAM devices to for delivering small β-cell clusters. As smaller cell clusters (100-150 μm) have been shown to be superior as compared to random-sized pancreatic islets (50-400 μm) (*1*), we estimated device dimensions to house 500,000 β-cell clusters with sizes of 100 μm and 125 μm. Our calculations assume that the volume of 1 IEQ is 1.77 nL (*2*), while the volume of a 100 μm cell cluster is 0.52 nL and that of a 125 μm cluster is 1.02 nL. This results in a conversion of 60,000 IEQ/mL to 204,230 clusters/mL for 100-μm clusters and 104,117 clusters/mL for 125-μm clusters to achieve an equivalent volume faction. (**A**) A proposed scaled-up device designed to encapsulate 500,000 clusters of 100 μm consists of five cell pouches, each 50 mm in length and 5 mm in diameter, with a capacity of 0.5 mL per pouch. These pouches are arranged in parallel. The final dimensions of the device, including the iEOG and connectors, are approximately 29 mm x 68 mm, about 40% of the size of a credit card. (**B**) A proposed scaled-up device to encapsulate 500,000 clusters of 125 μm consists of five cell pouches, each measuring 60 mm in length and 7.5 mm in diameter, with a capacity of 0.95 mL per pouch. These pouches are similarly arranged in parallel to accommodate the ~500,000 125 μm clusters. The device, including the iEOG and connectors, measures about 41.5 mm x 78 mm, which is roughly 70% of a credit card's size. For both designs, the distance (d) between cell pouches is 1 mm, and the thickness of the cell-laden alginate hydrogel layer (t) is 0.75 mm.

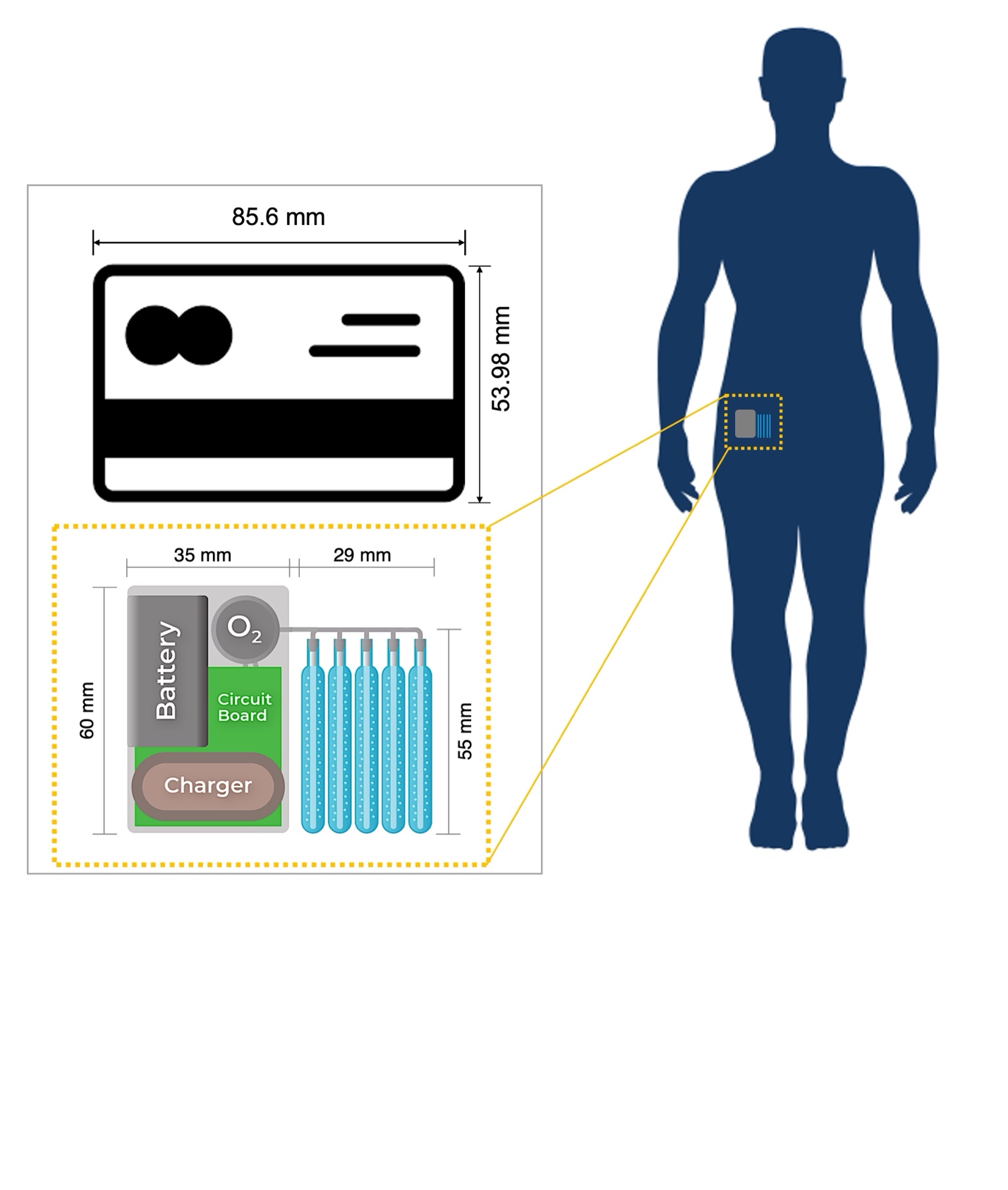

**Supplementary Figure 26**. A proposed design for a fully implantable BEAM system includes cell pouches encapsulating 500,000 clusters of 100 μm, a rechargeable battery, a transcutaneous energy transfer (TET) charger, and a circuit board. The entire system has a compact footprint of approximately 60 mm × 70 mm—roughly the size of a credit card. Additional batteries can be mounted on the reverse side of the board to extend operational time. A hydrogen diffusion membrane can be integrated into the surface of the electronics housing.

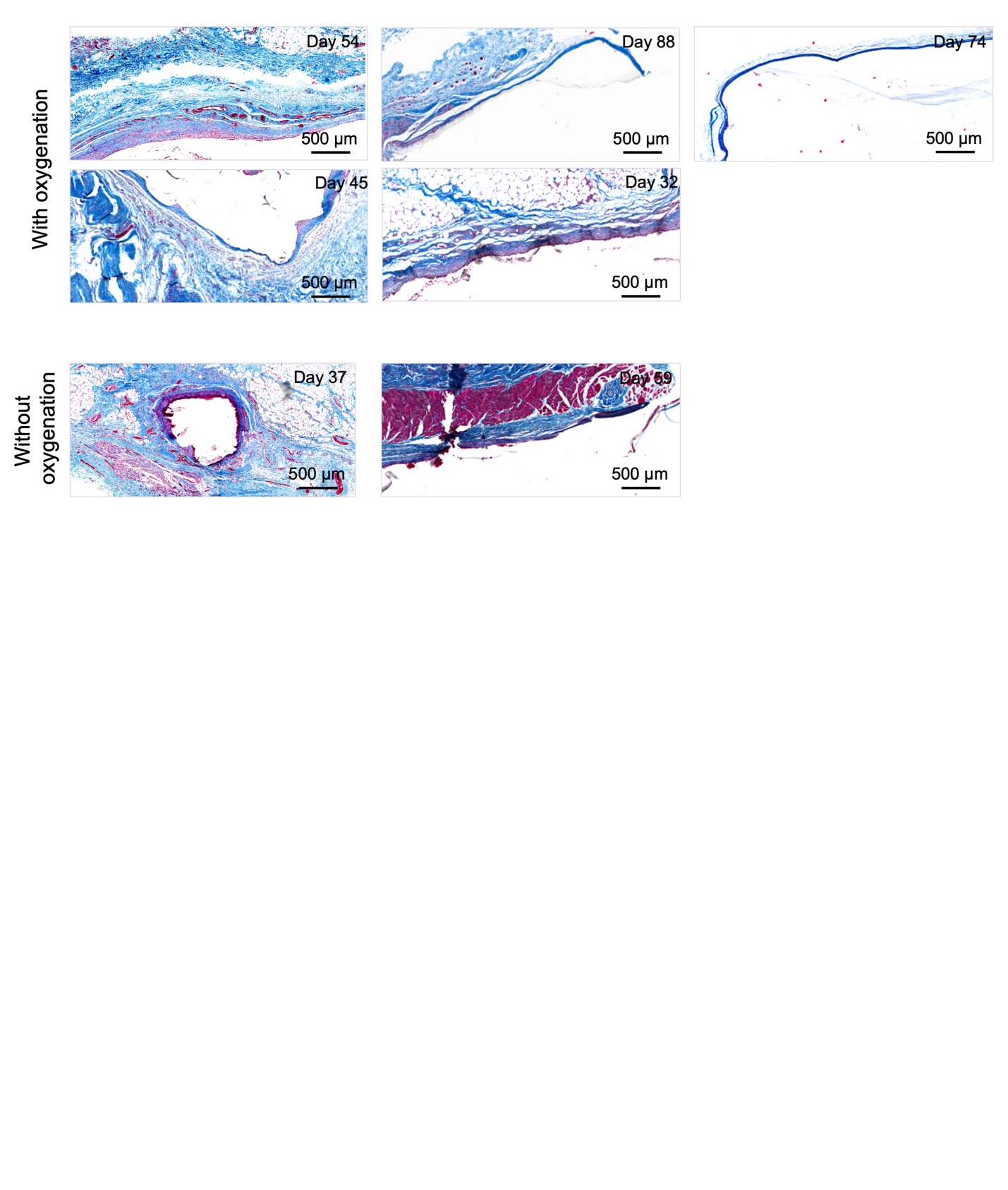

**Supplementary Figure 27**. Heterogeneity of foreign body responses and neovascularization around the devices with and without oxygen supplementation. Scale bar: 500 μm.

| Cell type | OCR (fmole/min/cell) | OCR (pmole/min/IEQ) | OCR (nmole/min/mg DNA) | Specific conditions | Ref. |
| --- | --- | --- | --- | --- | --- |
| INS-1 | ~2.6 | ~4.056 | ~390 | 2.5 mM glucose in DMEM | (*3*) |
|  | ~3.6 | ~5.616 | ~540 | 16.7 M glucose in secretion assay buffer | (*4*) |
|  | ~2.4 | ~3.744 | ~360 | 2.5 mM glucose in secretion assay buffer | (*4*) |
|  | ~3 | ~4.68 | ~450 | 3 mM glucose | (*5*) |
| Rat islets | 2.2 | 3.5 | 326.9231 |  | (*6*) |
|  | 3.82 ± 0.45 | 5.959 ± 0.702 | 572.981 ± 67.5 | DMEM containing 4.5 g/L glucose | (*7*) |
|  | 2.733 ± 0.867  2.4 ± 0.733  2.267 ± 0.4 | 4.264 ± 1.532  3.744 ± 1.144  3.536 ± 0.624 | 410 ± 130  360 ± 110  340 ± 60 | Day 0 after isolation  At 37^o^C  DMEM containing 4.5 g/1 glucose | (*8*) |
|  | 2.8 ± 0.867  3 ± 0.867  2.867 ± 0.8 | 4.368 ± 1.352  4.68 ± 1.352  4.472 ± 1.248 | 420 ± 130  450 ± 130  430 ± 120 | Day 2 after isolation at 37^o^C  DMEM containing 4.5 g/1 glucose |  |
| Human islets | 0.796 | 1.24176 | 119.4 |  | (*9*) |
|  | 0.92 ± 0.36 | 1.4352 ± 0.5616 | 138 ± 54 | CMRL 1066 with 10% FBS and 1% heparin | (*10*) |
|  | 1.3467 ± 0.58 | 2.1008 ± 0.9048 | 202 ± 87 | CMRL 1066 with 10% FBS and 1% heparin | (*10*) |
|  | ~0.9333 | ~1.456 | ~140 |  | (*11*) |
|  | 0.767 ±  0.733 ± | 1.196 ± 0.052  1.144 ± 0.062 | 115 ± 5  110 ± 6 | CMRL 1066 with 10% FBS and 1% heparin | (*12*) |
|  | 1.533 ± | 2.392 ± 0.624 | 230 ± 60 | Day 0 after isolation at 37^o^C  DMEM containing 4.5 g/1 glucose | (*8*) |
|  | 1.867 ± | 2.912 ± 1.144 | 280 ± 110 | Day 2 after isolation at 37^o^C  DMEM containing 4.5 g/1 glucose | (*8*) |
| Non-human primate islets | 0.5135 | 0.80106 | 77.025 | Basal glucose concentration in CMRL 1066  (24 h) | (*13*) |
|  | 0.2902 | 0.45267 | 43.526 | Basal glucose concentration in CMRL 1066  (168 h) |  |
| SC-islets | ~0.8 | ~1.248 | ~120 |  | (*11*) |
| SC-β cells | 1.506 | 2.34936 | 225.9 |  | (*1*) |

**Supplementary Table 1**. **Oxygen consumption rates of insulin-secreting cells**. For ease of comparison across studies employing varying units, the original values are presented in orange, while estimated values in alternative units are displayed in blue. The conversion is based on the assumption that 1 IEQ corresponds to 1,560 cells, containing 10.4 ng of DNA.

**Supplementary Table 2**. Summarization of incidents that happened during the animal study.

|  | Rat ID | Diabetes correction | Incidents |
| --- | --- | --- | --- |
| Design 1 | Rat 1 | ✓ | Transcutaneous oxygen tubing was damaged by the rat on day 3. |
|  | Rat 2 | ✓ | Transcutaneous oxygen tubing was damaged by the rat on day 1. |
|  | Rat 3 | ✓ | Transcutaneous oxygen tubing was damaged by the rat on day 5. |
| Design 2  (Transcutaneous tubing was hidden under the jacket) | Rat 4 | ✓ | Gas leakage from transcutaneous tubing on day 51-54. |
|  | Rat 5 | ✓ | Short iEOG disconnection on day 79 and 88 due to rat escaping from jacket. The device was retrieved ~ 2 h after the second disconnection. A small gas bubble was found around the cell pouch. |
|  | Rat 6 | ✓ | Short iEOG disconnection on day 67 due to rat escaping from jacket. iEOG was disconnected for more than 24 h on day 71. Gas bubbles were not found around the cell pouch. |
|  | Rat 7 | ✓ | No disconnection or device malfunction. A gas bubble was found around the cell pouch. |
|  | Rat 8 | ✓ | No disconnection or device malfunction. A gas bubble was found around the cell pouch. |
|  | Rat 9 | ✓ | No disconnection or device malfunction. A gas bubble was found around the cell pouch. |

**Supplementary Table 3**. Loading densities employed in micro- and macro-encapsulation systems. The summary is restricted to systems utilizing primary islets or SC-β cells that demonstrate diabetes reversal for a duration exceeding two weeks in immunocompetent animal models.

|  | Type of encapsulation | Loading density | Cell type | Ref. |
| --- | --- | --- | --- | --- |
| Without oxygenation | Microencapsulation | 1000 islets/mL | Rat islets | (*14, 15*) |
|  | Microencapsulation | 1538 IEQ/mL | Rat islets | (*16*) |
|  | Microencapsulation | 4000-16,000 IEQ/mL | Human islets | (*17*) |
|  | Microencapsulation | 2000 cell clusters/mL | SC-beta cells | (*18*) |
|  | Microencapsulation | 4000 IEQ/mL | Porcine islets | (*19*) |
|  | Microencapsulation | 200-2000 cell clusters/mL | SC-beta cells | (*20*) |
|  | Macroencapsulation | 25,000 IEQ/mL | Rat islets | (*21*) |
|  | Macroencapsulation | 8300 IEQ/mL | Rat and human islets | (*22*) |
|  | Macroencapsulation | 25,000 islets/mL | Rat islets | (*23*) |
|  | Macroencapsulation | 25,000 IEQ/mL | Rat islets | (*24*) |
| With oxygenation | Macroencapsulation | 2250 IEQ/cm^2^ | Rat islets | (*25*) |
|  | Macroencapsulation | 1000-4800 IEQ/cm^2^ | Rat islets | (*26*) |
|  | Macroencapsulation | 6250-8330 IEQ/mL | Rat islets | (*27*) |
|  | Macroencapsulation | 1000 IEQ/cm^2^ or 21,250 islets/mL | Rat islets | (*28*) |
|  | Macroencapsulation | 60,000 IEQ/mL or 4200 IEQ/cm^2^ | Rat islets | This paper |

**Supplementary Table 4**. The capacity estimation and the corresponding cell doses that can be loaded into a cell encapsulation pouch of the BEAM device.

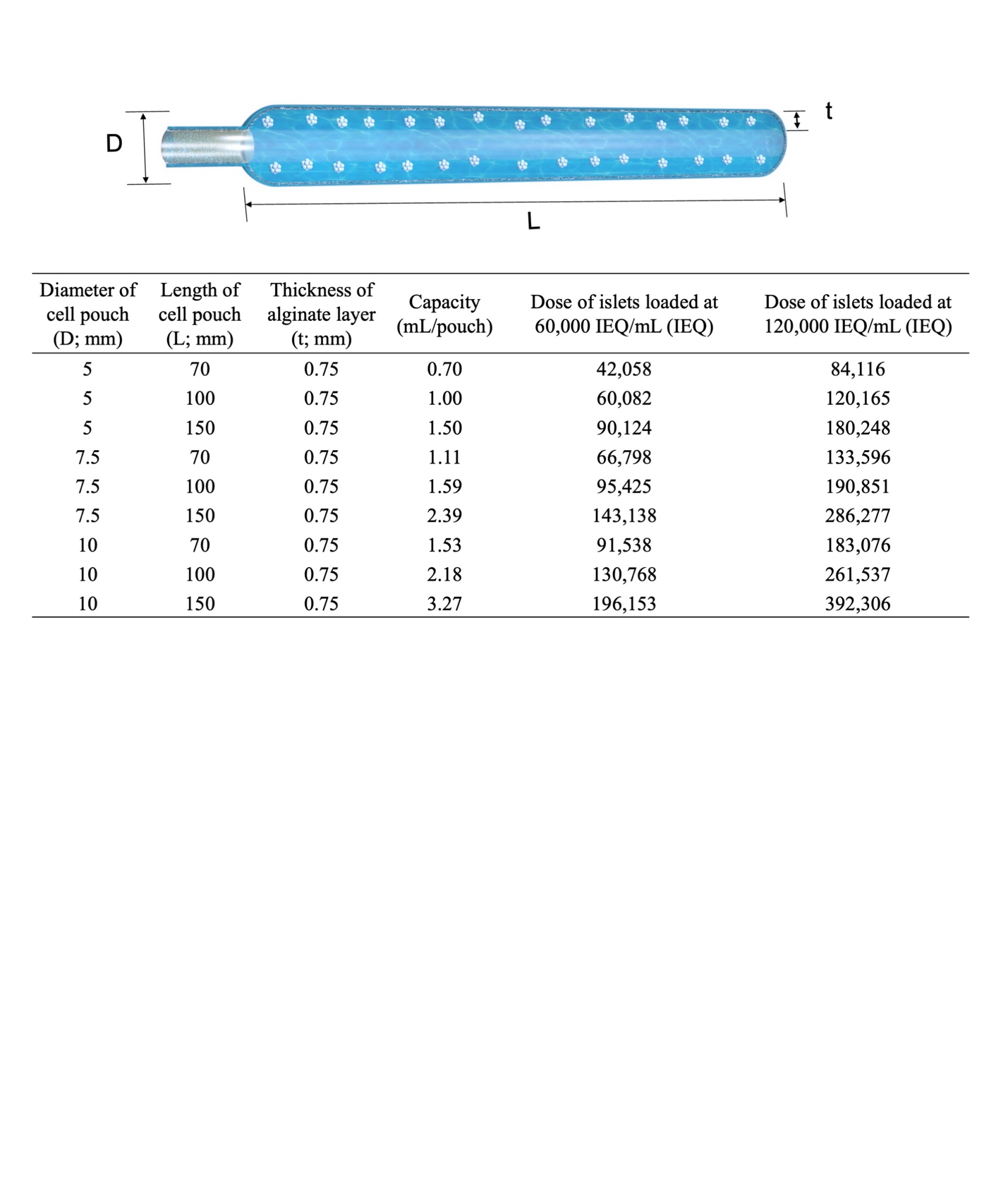
